# Identification, Isolation, and Characterization of Human LGR5-positive Colon Adenoma Cells

**DOI:** 10.1101/118034

**Authors:** Michael K. Dame, Durga Attili, Shannon D. McClintock, Priya H. Dedhia, Peter Ouilette, Olaf Hardt, Alana M Chin, Xiang Xue, Julie Laliberte, Erica L Katz, Gina M. Newsome, David R. Hill, Alyssa J. Miller, Yu-Hwai Tsai, David Agorku, Christopher H Altheim, Andreas Bosio, Becky Simon, Linda C Samuelson, Jay A Stoerker, Henry D Appelman, James Varani, Max S. Wicha, Dean E. Brenner, Yatrik M. Shah, Jason R Spence, Justin A. Colacino

## Abstract

The intestine is maintained by stem cells located at the base of crypts and distinguished by the expression of LGR5. Genetically engineered mouse models have provided a wealth of information about intestinal stem cells, while less is known about human intestinal stem cells due to difficulty detecting and isolating these cells. We established an organoid repository from patient-derived adenomas, adenocarcinomas, and normal colon, which we analyzed for variants in 71 colorectal cancer (CRC) associated genes. Normal and neoplastic colon tissue organoids were analyzed by immunohistochemistry and fluorescent-activated cell sorting for LGR5. LGR5-positive cells were isolated from 4 adenoma organoid lines and were subjected to RNA-sequencing. We found that LGR5 expression in the epithelium and stroma was associated with tumor stage, and by integrating functional experiments with LGR5-sorted cell RNA-seq data from adenoma and normal organoids, we found correlations between LGR5 and CRC-specific genes, including *DKK4* (dickkopf WNT signaling pathway inhibitor 4) and *SMOC2* (SPARC related modular calcium binding 2). Collectively, this work provides resources, methods and new markers to isolate and study stem cells in human tissue homeostasis and carcinogenesis.

## INTRODUCTION

In adult mammals, the intestine is a site of rapid cellular turnover, mediated by a population of intestinal stem cells (ISCs) that reside at the base of intestinal crypts (Barker et al., 2008). These stem cells are identified by expression of several genetic markers in the mouse (van der Flier et al., 2009, Montgomery et al., 2011, Powell et al., 2012, Yan et al., 2012), and genetically modified mice provide a robust toolbox for isolating and manipulating these ISCs (Barker et al., 2007, Snippert et al., 2010, Sato et al., 2009). Lgr*5* (leucine-rich repeat-containing g-protein coupled receptor 5) is one of the best characterized of these markers. The LGR family of proteins (LGR4, 5, 6) code for receptors of the secreted R-spondin proteins (Rspo1- 4). Together, Lgr/Rspo acts to potentiate Wnt pathway signaling (Carmon et al., 2012, de Lau et al., 2011). Lineage tracing experiments have demonstrated that differentiated cell lineages in mouse small intestine and colonic crypts are clonally derived from *Lgr5*(+) ISCs (Barker et al., 2007). Additionally, lineage tracing studies have revealed that Lgr5 marks a population of stem-like cells within precancerous adenoma tissue that drive adenoma growth (Schepers et al., 2012), and human colorectal cancers overexpress *LGR5* (Junttila et al., 2015). Previous efforts to expand, isolate, and experimentally characterize primary human LGR5 cells have been hampered by two distinct issues: (1) The difficulty in obtaining cultures highly enriched for epithelial stem cells (Wang et al., 2015b), and (2) a paucity of specific reagents to detect and isolate live LGR5(+) human cells (Barker, 2014). Recent efforts have successfully used gene editing techniques to create human organoid *LGR5* reporter lines (Shimokawa et al., 2017), however, this approach does not allow isolation from primary (unmodified) tissue and is not broadly useful across many cell lines. Previous studies have also reported varied localization of Lgr5 within the normal crypt using antibody-based methods (Becker et al., 2008, Kleist et al., 2011, Fan et al., 2010, Takahashi et al., 2011, Kobayashi et al., 2012, Kemper et al., 2012). Efforts have also utilized *in situ* RNA hybridization strategies to detect *LGR5*+ cells in human tissue, however, this approach does not permit isolation of cells (Jang et al., 2013, Baker et al., 2015). Collectively, these studies highlight the need for approaches that can be widely adopted to identify and isolate human intestinal stem cells.

Intestinal organoid culture is a robust system to grow tissue-derived cells that maintains the cellular heterogeneity in the intestinal epithelium (Sato et al., 2011a, Dedhia et al., 2016). Here, we establish an organoid biobank of normal, precancerous adenoma and colon adenocarcinoma from human tissue. We report detailed and robust methods to identify and isolate human colon LGR5(+) cells from normal human intestine and primary adenomas growing in long-term organoid culture using commercially available antibodies. Using these methods, we quantify Lgr5 protein expression in human intestinal tissues, including a colon adenocarcinoma tissue microarray (TMA). Using magnetic bead and fluorescent activated cell sorting (FACS) to enrich and isolate LGR5(+) and LGR5(−) cells from organoids, we conducted RNA sequencing and defined the expression profile of human LGR5(+) adenoma cells. We generated a *TCF/LEF*-GFP reporter adenoma organoid line to test the relationship between WNT reporter activity and Lgr5 expression. Through our analysis, we find that LGR5 protein expression is associated with progressive stages of cancer and we demonstrate that isolated human LGR5+ cells are enriched for *TCF/LEF*-GFP expression, *SMOC2*, and *DKK4*, a gene not detectable in normal colon tissue but associated with colorectal cancer. The methods and datasets presented are powerful resources for basic biological studies of the role of LGR5(+) cells in human colon homeostasis, as well as translational studies in chemoprevention and precision medicine designed to target LGR5(+) cell populations.

## RESULTS

### Isolation, Culture and genomic characterization of human adenoma organoids

We have developed an ongoing repository (Table 1) of organoids from patient-derived adenomas (n=17; including 2 high-risk sessile serrated adenomas), adenocarcinomas (n=4; including 1 colitis-associated cancer), and normal colon (n=9). All have been cryopreserved at early passage, reestablished from frozen stock, validated as matching their source tissue by short tandem repeat profiling and genomic sequencing (see below sequencing methods), are mycoplasma-free, and have demonstrated long-term culture (i.e., > 6 months in continuous culture). The neoplasm-derived organoids have been genomically characterized for variants in a panel of common CRC driver mutations across 71 genes (Table 1).

**Table 1.** Repository of patient-derived colon adenoma and adenocarcinoma organoids. A targeted colorectal cancer DNA sequencing panel was used to determine the presence of variants for 71 different oncogenes and tumor suppressor genes often mutated in colorectal cancers. Stop codon (*); frame shift (FS).

| Neoplasm | ID | Sex & Age |  | Location | Variants Detected |
| --- | --- | --- | --- | --- | --- |
| Adenoma (45mm) | 282 | F | 66 | ascending | APC Ala759fs, APC Gln1429*, ATM Arg337His, ATM Lys926Glu, EP300 Pro1802Pro, MSH2 R909Q |
| Adenoma (10mm) | 300 | M | 59 | transverse | APC Ser985fs, APC Arg1450*, BRCA1 Leu954Leu, BRAF Thr599dup, APC E1317Q |
| Adenoma (60mm) | 469 | M | 86 | cecum, ascending | APC Gln1378*, APC Glu536*, TCF7L2 Ser35fs, KRAS Gly12Asp, KRAS Gly12Ser |
| Adenoma (≥10mm) | 569 | F | 18 | sigmoid | APC Ser1355fs, ATM Phe858Leu, ATM Pro1054Arg, KRAS Ala146Thr, KRAS Gly12Val. |
| Adenoma (15mm) | 574 | F | 69 | hepatic flexure | APC Arg876*, APC Glu1464* |
| Adenoma (30mm) | 575 | M | 73 | ascending | APC Arg1450*, APC His864fs, DMD Lys34Asn, KRAS Gln61His |
| Adenoma (20mm) | 584 | M | 61 | ascending | APC Thr1556fs, KRAS Ala146Val, MET Arg988Cys, PMS2 Gly29Ala, TP53 Arg267Trp, EP300 Asp1579Asn |
| Adenoma (35mm) | 590 | F | 58 | ascending | BUB1B Arg550*, FLCN His429fs, MLH1 Lys443fs, MSH3 Lys381fs, PALB2 Met296fs, TCERG1 Arg889fs, CTNNA1 Met826Thr, CTNNB1 Ser45Phe, MAP2K4 Val127Ala, MLH3 Pro564Ser, PIK3R1 Arg188Cys |
| Adenoma (15mm) | 726 | M | 51 | ascending | APC Ser837*, APC Arg1450* and AXIN2 Phe791Cys |
| Adenoma (25mm) | 735 | M | 72 | ascending | APC Leu665*, APC Arg1450*, CTNNB1 Ala126Ser, KRAS Gly12Asp |
| Adenoma (20mm) | 772 | F | 79 | rectum | APC Arg216*, APC Glu225*, AKT1 Gly393Gly, PIK3R1 Ile82fs |
| Adenoma (adjacent to adenocarcinoma) | 14-881 | M | 46 | rectum | TP53 Pro151Ser, KRAS Gly12Asp, APC Pro1443fs |
| Adenoma: FAP (2 mm) | 236 | F | 26 | ascending | APC Thr1556fs, APC Leu143fs, MLH3 E624Q |
| Adenoma: Lynch | 610 | F | 59 | ascending | MSH3 Lys383fs, POLE Pro441Leu, DMD Val2498Ala, APC Ser1321Ser, KRAS Gln61His, PIK3CA Glu545Lys, ERBB2 Tyr411Cys, CDH1 His121Tyr, BLM Asn515fs, ERBB2 Pro1207His, MLH3 Asn674fs, APC Arg976fs, FBXW7 Ser668fs, AXIN2 Gly665fs, APC Ser1465fs, TCERG1 Arg958fs, BRCA2 Val1852Ile, GPC6 Gly243Trp, BRCA2 Lys1691fs, BRCA2 Thr3033fs, MUTYH Pro65fs, TCF7L2 Lys485fs, SLC9A9 Ala268Val, ERBB2 Arg47His, MSH2 R909Q |
| Adenoma (2-5mm) | 664 | M | 52 | cecum | MSH6 A20V |
| Adenoma: Sessile serrated (20mm) | 245 | F | 54 | ascending | BRAF Val600Glu, WBSCR17 Ser432Ser |
| Adenoma: Sessile serrated (23mm) | 708 | F | 76 | cecum | BRAF Val600Glu, DCC Asn472fs |
| Adenocarcinoma | 781 | F | 52 | descending | PIK3CA Glu545Lys, TP53 Phe113delPhe, APC Arg232* |
| Adenocarcinoma | 815 | F | 74 | cecum | KRAS Gly12Asp, APC Glu1309*, APC Glu1379*, MLH3 Asp1131Gly, EGFR Val323Ile, AKT1 Thr197Thr |
| Adenocarcinoma | 861 | F | 76 | ascending | SMAD4 Arg361His, PMS2 Thr9Ser, BRAF Val600Glu, APC Thr1556fs, APC Leu749fs, SLC9A9 Ala445Thr, BRCA1 Ser1517Pro, CDC27 Ala15Thr, WBSCR17 Ala577Val, PIK3CA Asn345Lys, DMD Arg550His, SLC9A9 Ile397fs, PMS2 M622I |
| Adenocarcinoma: IBD-associated | 21 | M | 57 | ascending | APC Gly1312*, TP53 Gly266Arg, PIK3CA Glu726Lys, KRAS Ala146Thr, BRAF Gly469Ala, PIK3CA Arg38Cys |
| Normal colon | 78, 80, 81, 83, 84, 85, 87, 88, 89 |  |  | F (ages 29, 33, 45, 49, 52, 56, 62), M (ages 21, 55); ascending |  |

### LGR5 immunohistochemical specificity in human colon and intestine

Two antibody clones for human Lgr5 were used in this study: a rabbit monoclonal antibody generated against a peptide sequence of LGR5 (clone STE-1-89-11.5) and a rat monoclonal antibody generated against a full-length LGR5 protein (clone 22H2.8). At the onset of this study these antibodies were in development by Miltenyi Biotec, but now both antibodies are commercially available (Miltenyi Biotec GmbH, Bergisch Gladbach, Germany; see Methods and Supplemental Methods). Details of their development have been presented previously (Agorku et al., 2014, Agorku et al., 2013), including the use of human *LGR4* and *LGR6* stable transfectants to demonstrate lack of cross-reactivity with these close homologues.

LGR5 immunohistochemical (IHC) expression was localized with clone STE-1-89-11.5 to the crypt base columnar (CBC) cells in normal formalin fixed paraffin embedded (FFPE) colon tissue (Fig. 1A^1^). At high magnification this staining pattern marked thin cells (Fig. 1A^2^), consistent with the morphology of CBC cells. From the same patient, an adenoma (found in the adjacent margins of an adenocarcinoma tissue resection, 10cm from the histologically normal tissue) showed intensified staining at the dysplastic crypt bases (Fig. 1A^3^) and sporadic focal staining throughout the more disorganized epithelial component. Interestingly, stromal staining was pronounced in this cancer-associated adenoma (Fig. 1A^3^). Supportive *LGR5 in situ* hybridization (ISH) staining was observed in the normal CBC cells (Fig. 1B top panel); in the dysplastic epithelium (Fig. 1B, arrow-1) and in the associated stroma (Fig. 1B, arrow-2).

**Fig. 1.**
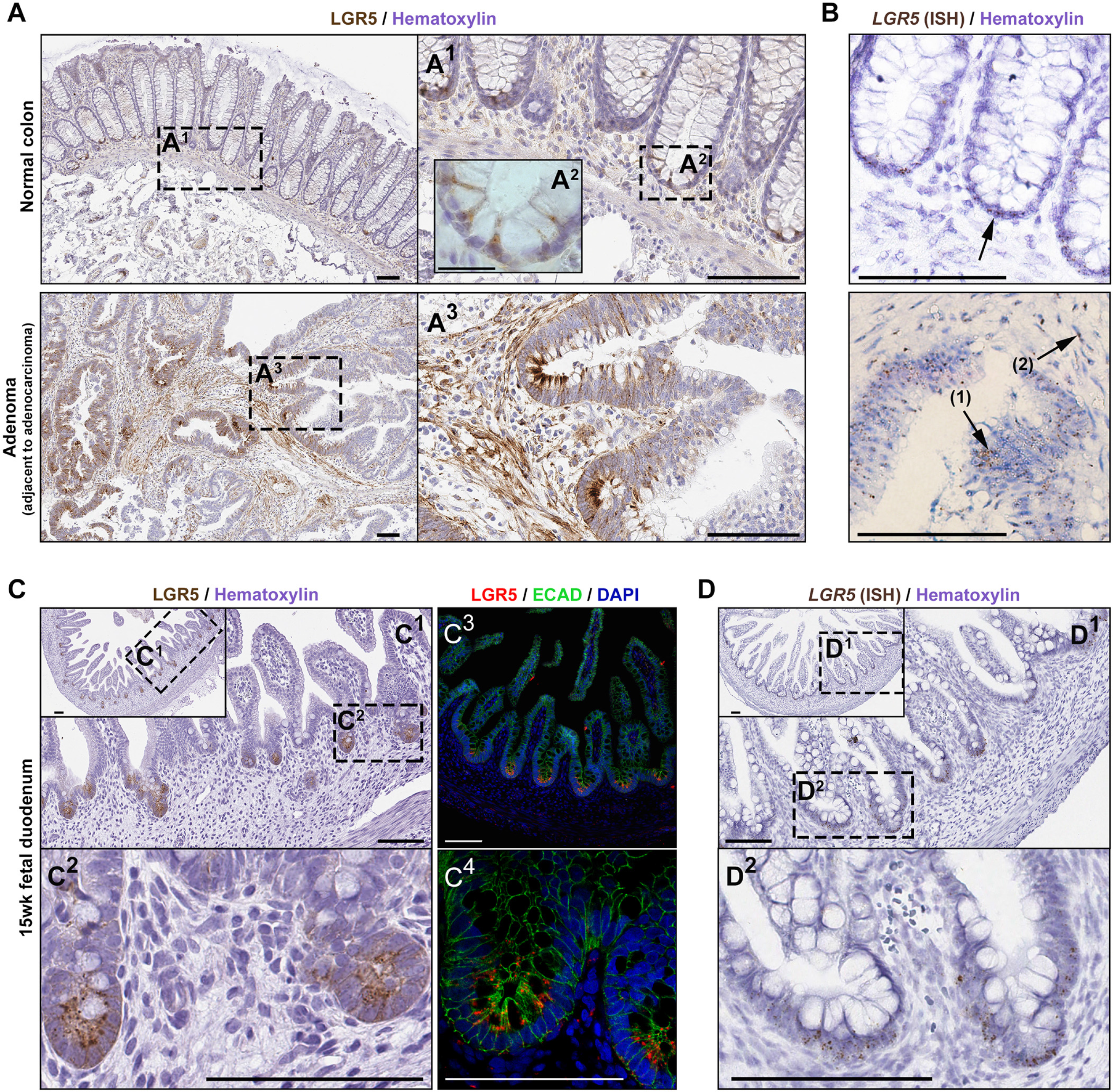
LGR5 immunochemical localization in human colon, colonic adenoma and duodenum. **(A)** LGR5 IHC staining in normal human colon (one of five representative patients) at low (A1) and high (A2) magnification, as well as adenoma (A3) from the same patient (high-grade dysplasia; adjacent to adenocarcinoma; specimen 14881). **(B)** *Lgr5* expression by *in situ* hybridization provides a conventional reference for the LGR5 IHC staining in normal crypts (upper panel) and in the adenoma (bottom panel); glandular *Lgr5* (arrow-1) and stromal expression (arrow-2) in adenoma; **(C)** LGR5 IHC (C1, C2) and IF staining (C3, C4) in fetal duodenum, and **(D)** ISH expression in the same duodenum specimen. Scale bars, 100μM; A2, 25μM.

The human fetal small intestine has been shown to express high levels of *LGR5* mRNA relative to adult by RNA-sequencing (Finkbeiner et al., 2015). Consistent with this, robust and specific Lgr5 protein staining by IHC and immunofluorescence (Fig. 1C), in conjunction with *LGR5* ISH (Fig. 1D), was observed in the proliferative zone of the 15-week fetal gut. In contrast, IHC and IF staining in adult duodenum (Fig. S1A) showed weak punctate LGR5(+) staining in cells present between Paneth cells marked by DefensinA5 (DEFA5), consistent with published ISH and RNA-sequencing data (Finkbeiner et al., 2015).

Clone STE-1-89-11.5 was further demonstrated to be specific for human LGR5 by Western blotting. Mouse 1881 lymphoma cells that were previously transfected with human *LGR5* served as a positive control [1881(+); provided by Miltenyi Biotec]. Transfection stability was confirmed by mRNA expression analysis (Fig. S1B). The antibody showed strong reactivity against the human LGR5 1881(+) cell line, as well as measurable activity against one adenoma organoid (specimen #14881) (Fig. S1C).

### LGR5 immunohistochemical expression levels, and localization, are associated with human colon cancer stage

LGR5 IHC staining was performed in a FFPE TMA, which included 2 normal tissues, 3 low grade small adenomas (well differentiated), and 70 adenocarcinomas. We also included staining of 5 additional normal colon autopsy samples obtained for our studies in this analysis (Fig. 2). Tissue sections were scored for staining intensity in the glandular epithelium and separately in the stroma. Unlike normal colon, which showed staining in CBCs at the base of the crypt, the adenomas stained intensely in distinct zones of epithelium or for the entire dysplastic crypt (Fig. 2A). Relative to normal tissue, the adenocarcinomas showed increased LGR5 staining in both the epithelium and in the stroma (Figure 2A-C).

**Fig. 2.**
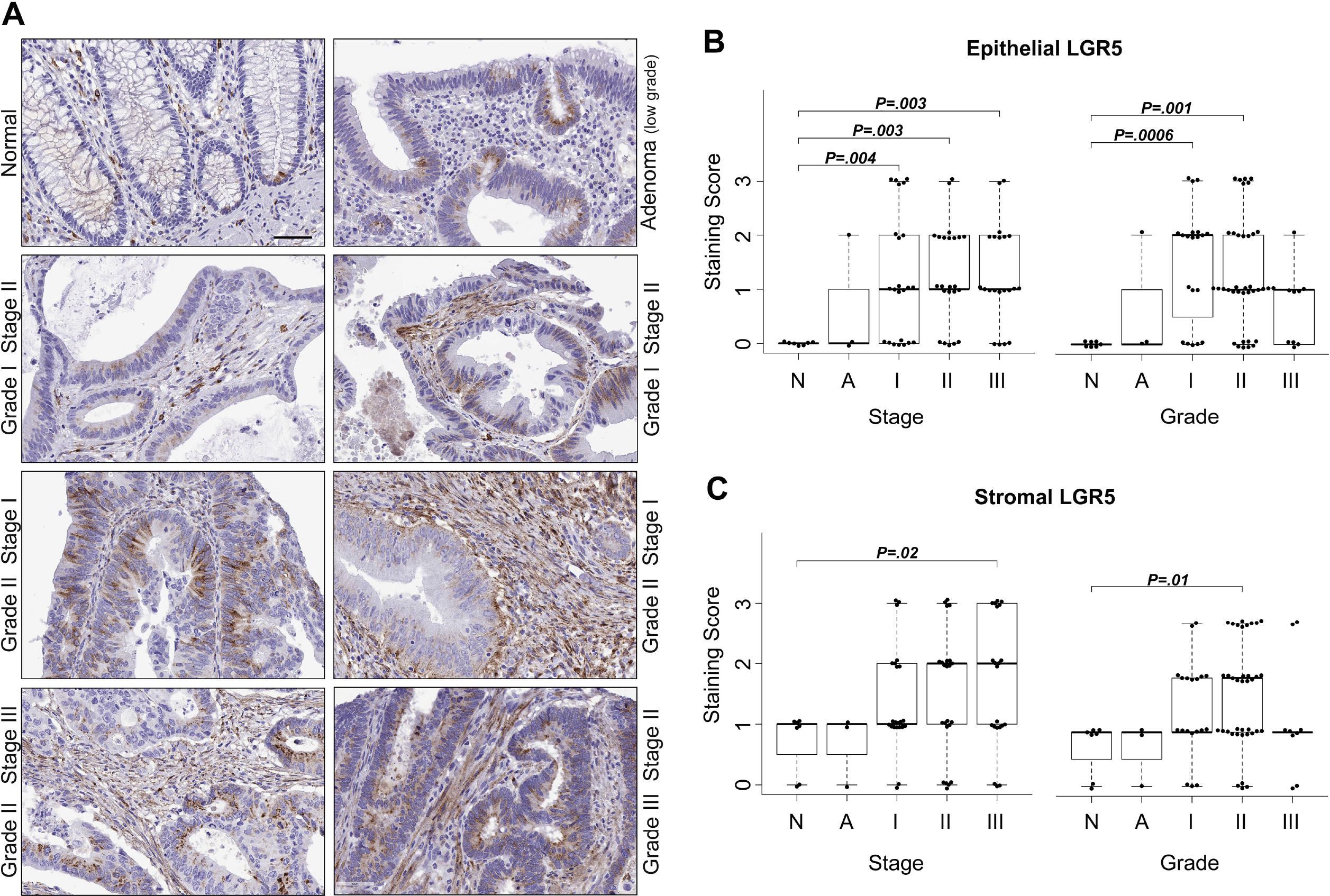
LGR5 immunohistochemical staining intensity in the epithelium and stroma across a range of colorectal adenocarcinoma stages and grades. **(A)** Representative images of LGR5 staining in the epithelial and stromal components in normal colon (n=5 normal autopsy samples) and from a CRC tissue microarray (n=2 normals, 65-68 neoplasm). Scale bar, 50μM. **(B)** Association between epithelial or **(C)** stromal staining intensity, and stage/grade of the tissue tested by generalized linear modeling, with staining intensity as the dependent variable and stage or grade as the independent variable.

### Isolating Lgr5(+) cells from patient-derived adenoma organoids using magnetically-activated cell sorting (MACS) and FACS

Fig. 3A outlines the approach developed for isolation and analysis of LGR5(+) cells from human adenoma. Adenoma organoids were established in culture (approx. 2 months) and genomically characterized (Table 1). The organoid cultures used in this study (14881, 282, 584, 590) were established and cultured in ‘reduced medium’, KGM-Gold™ (Lonza, Walkersville, MD; KGMG), a serum-free epithelial medium containing epidermal growth factor (EGF) and pituitary extract (Dame et al., 2014). We chose this medium to establish/expand non-normal epithelium, similar to approaches used in recent studies which have demonstrated that mutations allow cells to grow independent of particular ISC niche factors (Li et al., 2014, Matano et al., 2015, Drost et al., 2015). When organoid cultures grown in reduced medium were transferred to medium replete with ISC niche factors (L-WRN medium-see methods) (Miyoshi and Stappenbeck, 2013), they generally transitioned from budding structures (Fig. 3B, middle column) to thin-walled spherical cysts (Fig. 3B, right column). Accompanying this change in morphology was an increase in *Lgr5* mRNA expression in L-WRN-cultured organoids compared to organoids cultured in reduced medium, as assessed by qRT-PCR (Fig. 7A, left graph). Flow cytometric analysis revealed that L-WRN-cultured organoids also had an increase in the number of Lgr5 positive cells (Fig. 3C, comparing “stain” in the first and second row; Fig. S2A). Even with L-WRN-enhanced *LGR5* expression, a small proportion of cells were LGR5(+) based on flow cytometry, and therefore magnetic bead positive enrichment was used prior to FACS to increase the number of cells obtained.

**Fig. 3.**
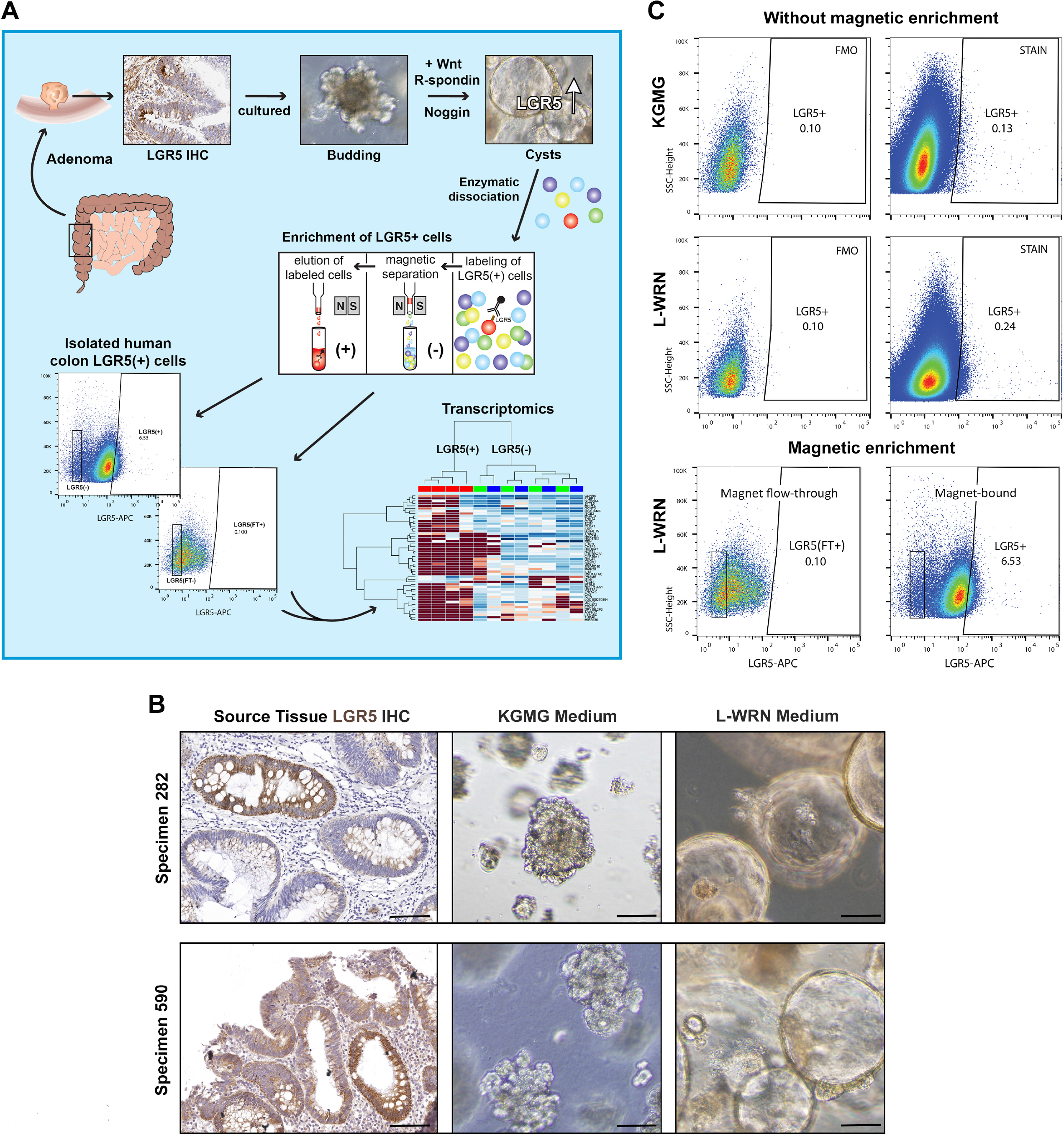
Strategy for the isolation of LGR5(+) cells from patient-derived colon adenoma. **(A)** Graphical representation of complete procedure. **(B)** Culture of adenoma organoids for LGR5 enrichment. LGR5 IHC staining (left column) in biopsied large adenoma (>10mm; two representative specimens are shown). Organoids with budding morphology (middle column) derived from these specimens cultured in the reduced medium KGMG. The same cultures with cystic morphology (right column) after being transferred for 3-4 weeks to L-WRN medium. Scale bars, 100μM. **(C)** Representative scatterplots of LGR5 expression in organoids cultured in KGMG or L-WRN (specimen 282). Live, LGR5(+) human adenoma cells were obtained after LGR5-magnetic bead enrichment by isolating DAPI(−) and LGR5(+) cellular populations.

Mouse 1881 cells stably expressing Lgr5 were used to validate the magnetic bead positive separation protocol and the high specificity of antibody clone 22H2.8. 1881-LGR5 expressing and nonexpressing cells were mixed in varying proportions, magnetically separated, and analyzed by flow cytometry (Fig. S2B). The initial ratio of 1881 LGR5(+) to 1881 LGR5(−) cells was predicted as shown by the pre-magnet flow histograms (Fig. S2B, left column). The post-magnetic separation shows almost complete separation of 1881 LGR5(+) cells in the positive fraction (Fig. S2B, right column) from 1881 LGR5(−) cells in the negative fraction (middle column). After magnetic bead based enrichment of the adenoma organoid cells, the unbound flow-through negative population was used to set the Lgr5 FACS gate (Fig. 3C bottom row). Magnetic bead-bound LGR5(+) cells from organoids were isolated via FACS, and importantly, we observed that while only a small percentage of cells bound to beads were LGR5(+) (Fig. 3C bottom row), on average, magnetic bead pulldowns of LGR5(+) cells led to an 11.3-fold enrichment in the number of LGR5(+) cells (cultured in L-WRN medium) obtained compared to samples without magnetic enrichment (specimens 282, 584, 590; SE=6.55; n=3 sorts no magnet, n=7 sorts with magnet; Table S1).

### Transcriptomic profiling of LGR5(+) adenoma cells

The gene expression signature of human LGR5(+) intestinal cells is essentially uncharacterized, which has significantly limited our understanding of the role of these cells in human intestinal stem cell biology and colorectal cancer progression. Using the methods described above, we isolated LGR5(+) cells from four different adenoma-derived organoid lines grown in L-WRN media and conducted RNA-seq on three sorted cellular populations from each (Fig. 4, Fig. S3): (1) magnetic column flow-through cells, termed LGR5(FT-); (2) cells that bound to the magnet, but were Lgr5 negative based on FACS, termed LGR5(−); and (3) cells that bound to the magnet and were LGR5 positive based on FACS, termed LGR5(+). Genomic variant analysis showed that each of these patient specimens has functionally significant mutations within commonly mutated colon cancer genes, confirming the adenoma, and not normal, identity of these organoids (Table 1). Expression profiles of the LGR5(−) and LGR5(FT-) were found to be largely similar at both the gene expression and pathway level (Fig. S3C, D). Due to transcriptional similarity between negative populations, our analysis focused on comparisons between the magnetic bead-bound cells deemed to be LGR5(+) and LGR5(−) by FACS. Multidimensional scaling on the top 500 most variably expressed genes revealed that samples clustered distinctly by the patient of origin, not by the expression of Lgr5 (Fig. 4A, Fig. S3A). Despite this, we identified 519 differentially expressed genes (FDR *P*-value < .05) between the two cell populations across four genetically diverse adenomas (Fig. 4B and Table S2). Expression values for all differentially expressed genes for each specimen, as well comparisons between LGR5(+) cells and LGR5(FT-) cells, are reported in Table S2. We determined that LGR5 had the highest level of statistical enrichment in the LGR5(+) cells (FDR=3.8E- 21) and was expressed an average of 5.5-fold higher compared to LGR5(−) cells, lending confidence to the specificity of the LGR5 antibody and the separation procedure. Unsupervised hierarchical clustering analysis using the top 50 most differentially expressed genes showed clear separation between the LGR5(+) and LGR5(−) samples from three of the four enteroid lines (Fig. 4C), suggesting both commonalities and heterogeneity in the LGR5(+) cell gene expression signature between specimens. When the expression signature of LGR5(+) cells were clustered with those from both the LGR5(−) and LGR5(FT-) cells, however, there was clear separation of the LGR5(+) cells across all four organoid lines (Fig. S3B). Stem cell markers associated with the colon as well as other tissue-specific stem cells including *BMI1, MEX3A*, and *SMOC2*, were upregulated in LGR5(+) cells, while known markers of colonic differentiation, including *MUC2, TFF3*, and *KRT20*, were down-regulated (Fig. 4D). KEGG pathway analyses (Kyoto Encyclopedia of Genes and Genomes) (Kanehisa and Goto, 2000) revealed differential expression in LGR5(+) cells for genes involved in metabolism, including the pathways, *glycolysis and gluconeogenesis, biosynthesis of amino acids, metabolic pathways, carbon metabolism, fructose/mannose metabolism, glycosphingolipid biosynthesis*, and *central carbon metabolism in cancer* (Fig. 4E; Fig. S3C). Additionally, LGR5(+) cells had significantly differentially expressed genes involved in *HIF-1 signaling, cytokine-cytokine receptor interaction*, *hematopoietic cell lineage*, *MAPK signaling*, *PI3K/AKT signaling*, and *pathways in cancer* (Table S2; Fig. 4E, Fig. S3C, Fig. S4-9).

**Fig. 4.**
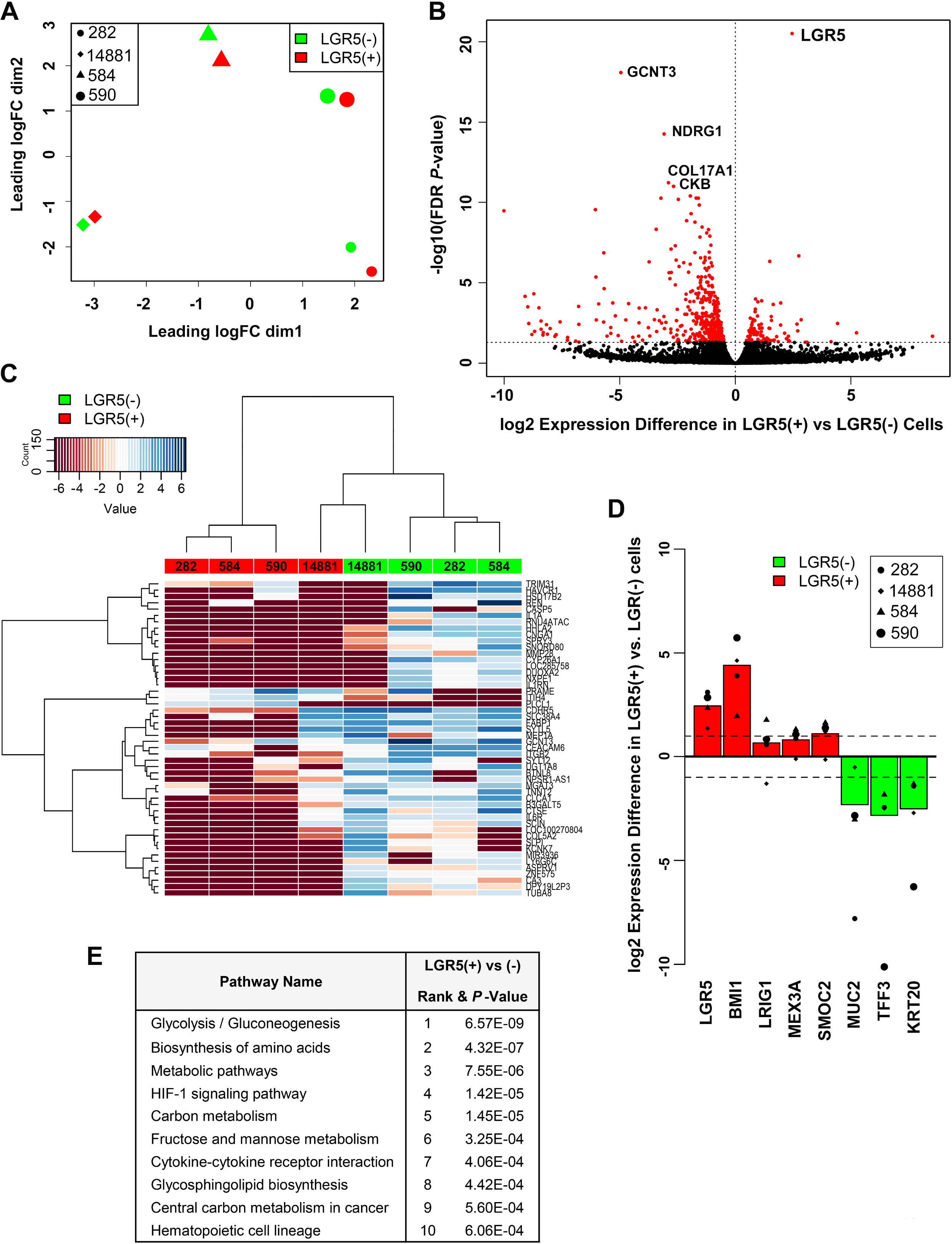
Transcriptomic profiling of human LGR5(+) and Lgr5(−) primary adenoma cells isolated from four patient-derived organoid cultures. **(A)** Multidimensional scaling plot of the LGR5(+) and LGR5(−) cells, based on the top 500 most variable genes. Patient identifiers #14881, 228, 584, and 590. **(B)** False discovery rate (FDR) volcano plot of the log(2) ratio of gene expression between the LGR5(+) and LGR5(−) cells. **(C)** Unsupervised hierarchical clustering heatmap of the 50 most differentially expressed genes by fold change between the LGR5(+) (red) and LGR5(−) (green) populations. **(D)** Log(2) fold change in gene expression between LGR5(+) and LGR5(−) cells for known markers of colon stem (red) and differentiated (green) cells. **(E)** The top 10 most enriched KEGG pathways for differentially expressed genes between the LGR5(+) and Lgr5(−) cells.

We also identified a high level of concordance between LGR5(+) cells and a previously reported set of genes associated with high levels of Wnt signaling in mouse cancer stem cells (based on TOP-GFP Wnt reporter activity (Vermeulen et al., 2010) (Fig. 5A). To functionally validate this overlap, we transduced organoids (line 282) with a lentiviral *TCF/LEF*-GFP reporter (Figure 5B). *TCF/LEF*-GFP organoids were stained with the Lgr5 antibody for analysis by ImageStream flow cytometry. ImageStream analysis visually established the specificity of the fluorescently tagged Lgr5 antibody and *TCF/LEF*-GFP expression to viable cells (Fig. 5C). Based on the same experimental paradigm established by Vermeulen et al., we compared LGR5 expression in the top 10% highest and lowest GFP expressing cells. We identified a significant enrichment of LGR5(+) cells in the *TCF/LEF*-GFP^HI^ cells relative to low cells. (average 3.8 fold enrichment of Lgr5; 6.2% vs. 1.6%).

**Fig. 5.**
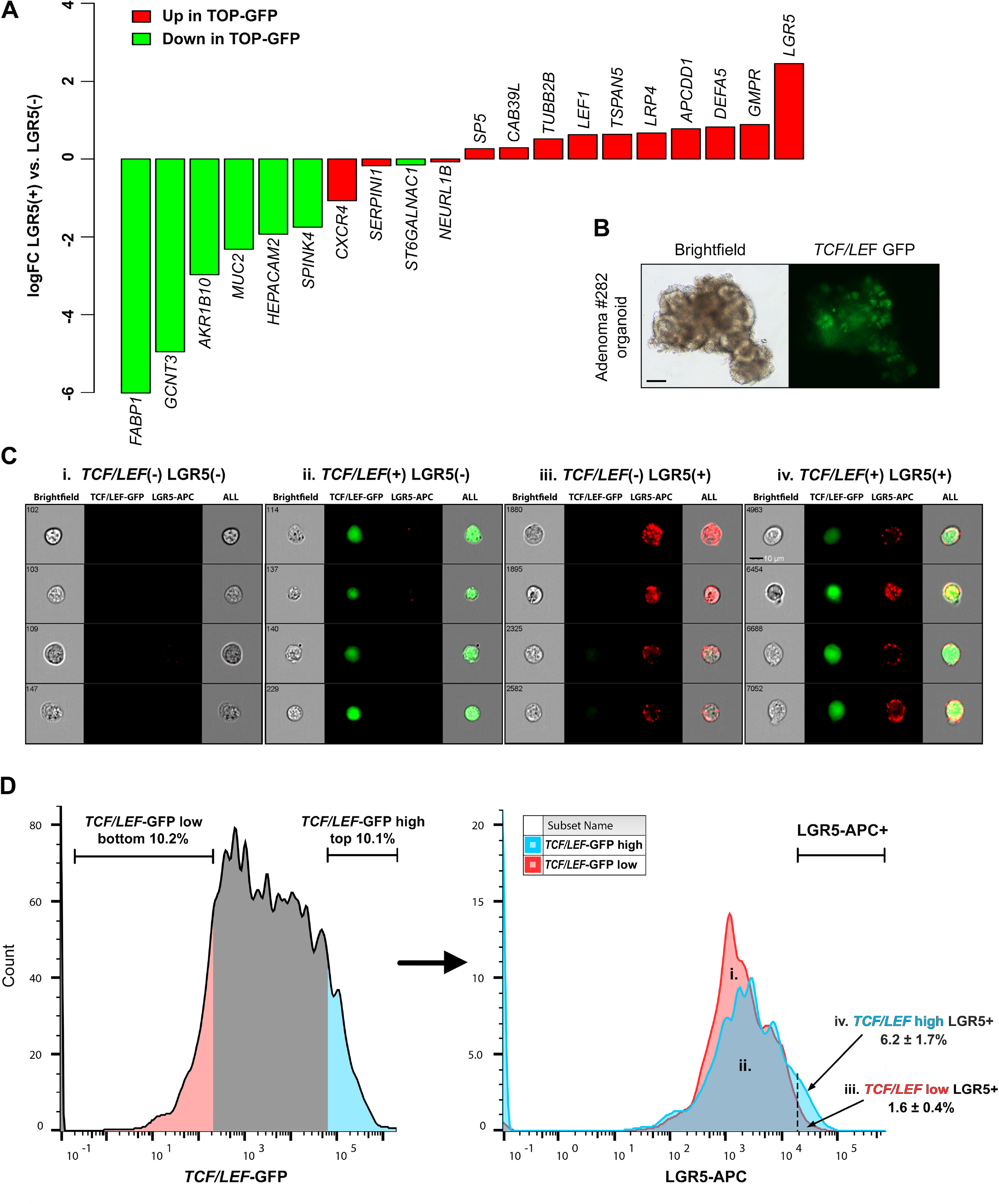
Comparisons and functional validation of the association between the human adenoma LGR5(+) cell gene signature and previously published expression signature in high-Wnt colon cancer stem cells. **(A)** Concordance of adenoma LGR5(+) cell gene expression with gene expression associated with Wnt signaling in colon cancer stem cells, based on TOP-GFP Wnt reporter activity (Vermeulen et al., 2010). **(B)** Brightfield and fluorescence images of the 282 adenoma organoid line lentivirally transduced with a *TCF/LEF*-GFP reporter for Wnt signaling. Scale bar, 50μM. **(C)** Representative ImageStream flow cytometry results of *TCF/LEF*-GFP organoids stained with the LGR5- APC antibody (n=3 experimental replicates). Scale bar, 10μM. (D) Comparison of LGR5 expression in the 10% highest and 10% lowest *TCF/LEF*-GFP expressing cells; values in iii and iv represent mean % ± s.e.m.

To identify a “human adenoma LGR5(+) cell gene signature” with more stringent parameters, we compared genes upregulated in LGR5(+) cells to both LGR5(−) and to LGR5(FT-) cells to which we identified 33 genes (Table 2). A comparison between these 33 genes in the “human adenoma LGR5(+) cell gene signature” and the previously published “murine intestinal stem cell signature,” generated from isolated *Lgr5*^HI^ ISCs (Munoz et al., 2012), identified 4 overlapping genes that were enriched in both populations, including the canonical ISC markers *LGR5, SMOC2*, and *CDCA7* (Fig. 6A). This analysis also revealed a large number of genes in the human adenoma LGR5(+) cell signature that did not overlap with the mouse *Lgr5* stem cell signature.

**Fig. 6.**
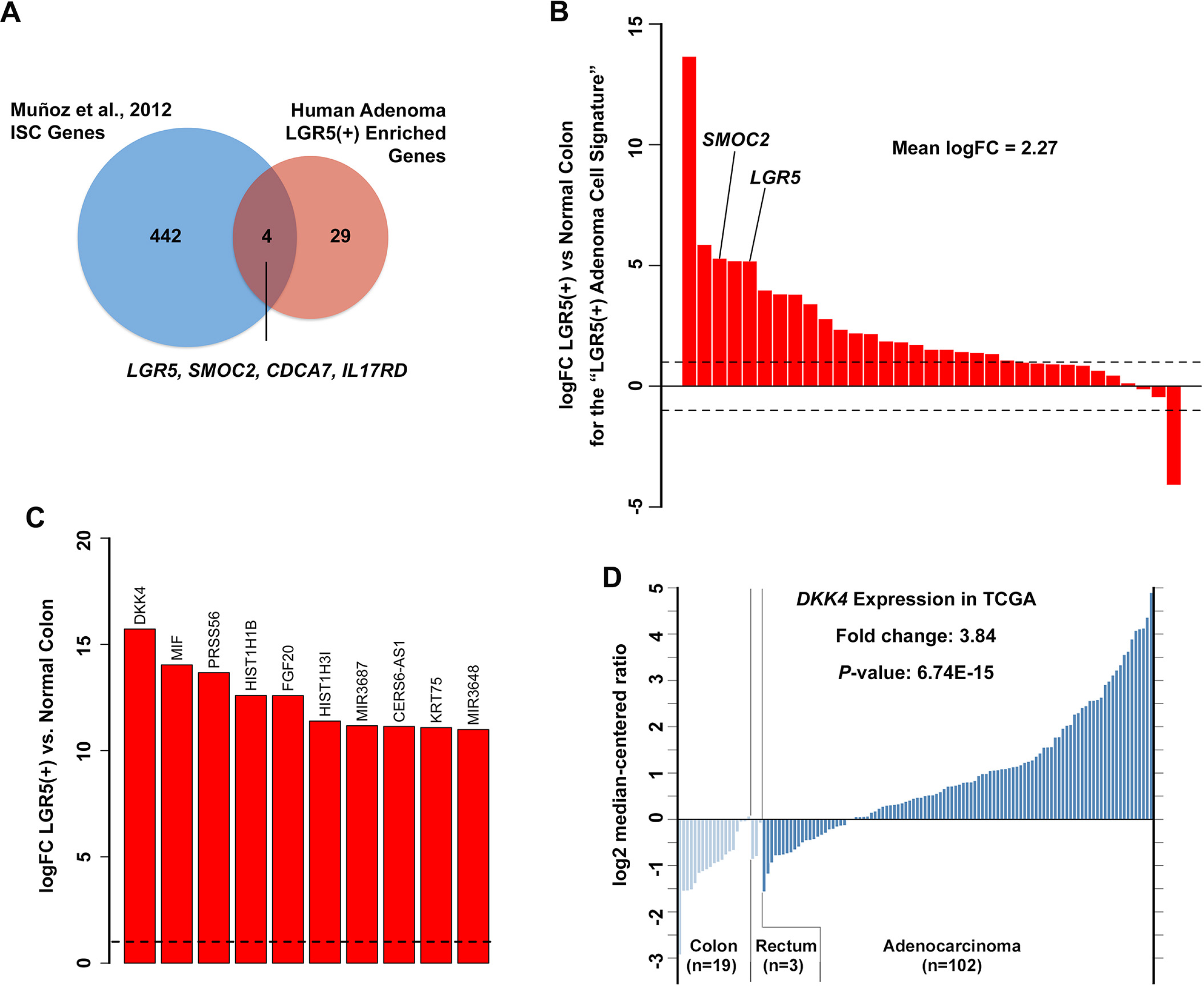
Comparisons between genes in the human adenoma LGR5(+) cell gene signature and the previously published murine intestinal stem cell signature, and the normal human colon signatures. **(A)** Gene expression overlap between genes upregulated in LGR5(+) vs. LGR5(−) cells and the *Lgr5* mouse intestinal stem cell signature reported in Munoz et al (Munoz et al., 2012). **(B)** Comparison of gene expression between LGR5(+) adenoma cells and previously published RNA-seq data from normal human colon (Uhlen et al., 2015) for genes in the “human adenoma LGR5+ cell gene signature. **(C)** The top ten most differentially expressed genes by magnitude between LGR5(+) adenoma cells and normal human colon (all genes; unbiased analysis) (Uhlen et al., 2015). **(D)** An analysis of TCGA colorectal cancer gene expression data through Oncomine™ of DKK4, comparing colorectal adenocarcinoma expression with normal colon and rectum.

**Table 2.**
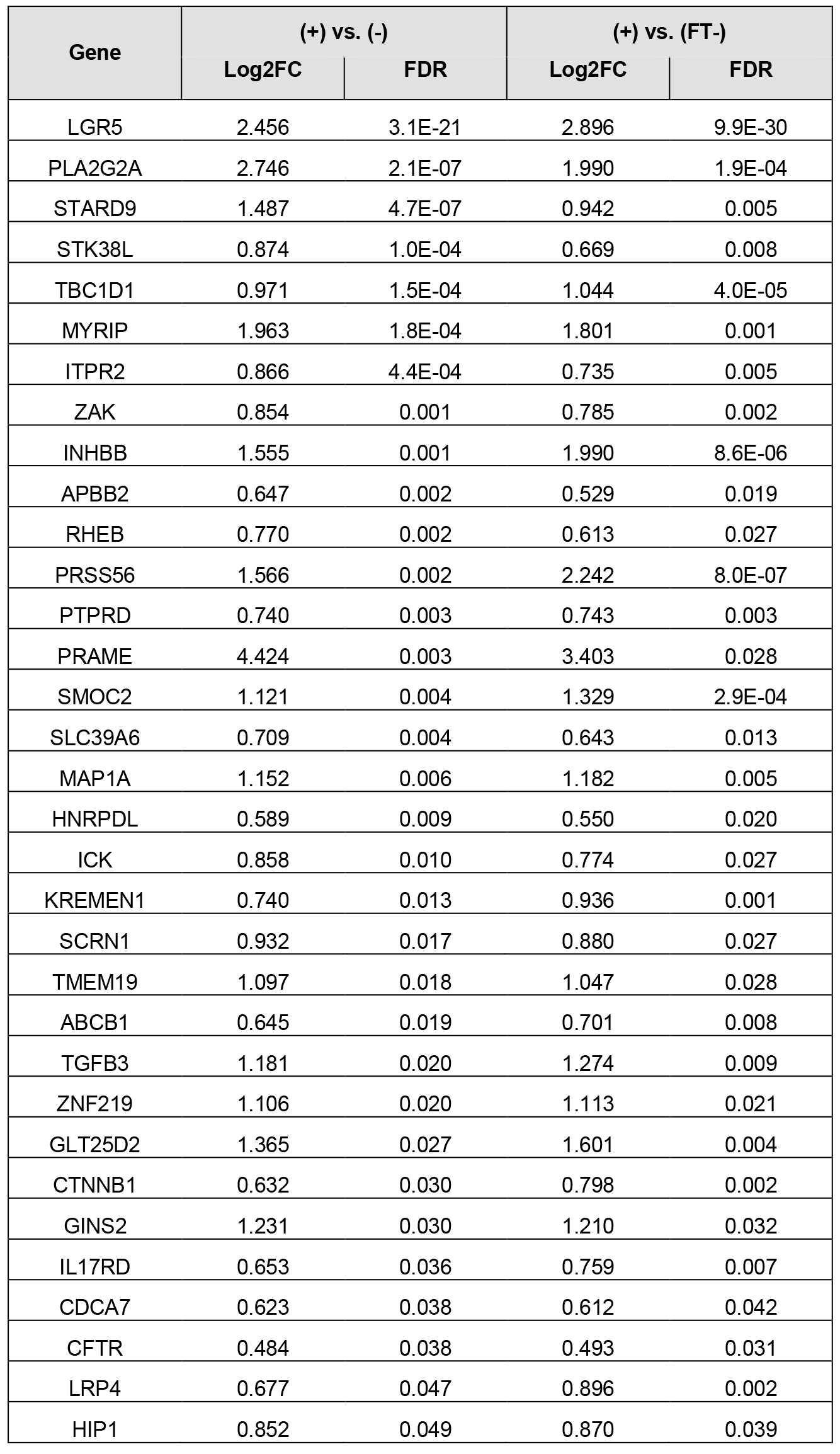
Genes upregulated in human adenoma LGR5(+) cells when compared to both Lgr5(−) and to LGR5(FT-) cells.

| Gene | (+ vs. (-) |  | (+ vs. (FT-)) |  |
| --- | --- | --- | --- | --- |
|  | Log2FC | FDR | Log2FC | FDR |
| LGR5 | 2.456 | 3.1E-21 | 2.896 | 9.9E-30 |
| PLA2G2A | 2.746 | 2.1E-07 | 1.990 | 1.9E-04 |
| STARD9 | 1.487 | 4.7E-07 | 0.942 | 0.005 |
| STK38L | 0.874 | 1.0E-04 | 0.669 | 0.008 |
| TBC1D1 | 0.971 | 1.5E-04 | 1.044 | 4.0E-05 |
| MYRIP | 1.963 | 1.8E-04 | 1.801 | 0.001 |
| ITPR2 | 0.866 | 4.4E-04 | 0.735 | 0.005 |
| ZAK | 0.854 | 0.001 | 0.785 | 0.002 |
| INHBB | 1.555 | 0.001 | 1.990 | 8.6E-06 |
| APBB2 | 0.647 | 0.002 | 0.529 | 0.019 |
| RHEB | 0.770 | 0.002 | 0.613 | 0.027 |
| PRSS56 | 1.566 | 0.002 | 2.242 | 8.0E-07 |
| PTPRD | 0.740 | 0.003 | 0.743 | 0.003 |
| PRAME | 4.424 | 0.003 | 3.403 | 0.028 |
| SMOC2 | 1.121 | 0.004 | 1.329 | 2.9E-04 |
| SLC39A6 | 0.709 | 0.004 | 0.643 | 0.013 |
| MAP1A | 1.152 | 0.006 | 1.182 | 0.005 |
| HNRPDL | 0.589 | 0.009 | 0.550 | 0.020 |
| ICK | 0.858 | 0.010 | 0.774 | 0.027 |
| KREMEN1 | 0.740 | 0.013 | 0.936 | 0.001 |
| SCRN1 | 0.932 | 0.017 | 0.880 | 0.027 |
| TMEM19 | 1.097 | 0.018 | 1.047 | 0.028 |
| ABCB1 | 0.645 | 0.019 | 0.701 | 0.008 |
| TGFB3 | 1.181 | 0.020 | 1.274 | 0.009 |
| ZNF219 | 1.106 | 0.020 | 1.113 | 0.021 |
| GLT25D2 | 1.365 | 0.027 | 1.601 | 0.004 |
| CTNNB1 | 0.632 | 0.030 | 0.798 | 0.002 |
| GINS2 | 1.231 | 0.030 | 1.210 | 0.032 |
| IL17RD | 0.653 | 0.036 | 0.759 | 0.007 |
| CDCA7 | 0.623 | 0.038 | 0.612 | 0.042 |
| CFTR | 0.484 | 0.038 | 0.493 | 0.031 |
| LRP4 | 0.677 | 0.047 | 0.896 | 0.002 |
| HIP1 | 0.852 | 0.049 | 0.870 | 0.039 |

Given that our transcriptional analysis compared cells, LGR5(+) vs. LGR5(−), isolated from adenoma-derived organoid cultures (as opposed to intact tissue biopsies), we reasoned that it was possible that LGR5(+) enriched genes might be masked by comparing fractions which, in whole, were cultured in medium that promotes high levels of WNT signaling (L-WRN). Therefore, we also compared LGR5(+) cells with previously published RNA-seq data from whole-thickness normal human colon (Uhlen et al., 2015) (Fig. 6). When we compared the expression of normal colon genes with the 33 genes from the “human adenoma LGR5(+) cell gene signature” (Table 2), we observed a strong enrichment for a majority of the 33 genes (Fig. 6B), including the *Lgr5* surrogate marker gene *SMOC2*. In an unbiased comparison of all LGR5(+) expressed genes, one of the most upregulated in LGR5(+) adenoma cells compared to normal colon was the Wnt pathway inhibitor *DKK4* (dickkopf WNT signaling pathway inhibitor 4) (Fig. 6C). *DKK4* was virtually undetectable in the normal colon (Table S2). An analysis of TCGA colorectal cancer gene expression data (The Cancer Genome Atlas, 2012) revealed that *DKK4* is significantly upregulated in colorectal tumors compared to normal tissue (Fig. 6D). To follow up on these correlations between *LGR5, SMOC2* and *DKK4* identified via RNA-seq analysis, we performed additional functional and expression profiling experiments. Three adenoma organoid lines were grown in reduced medium or in L-WRN media to drive a more “stem-like” phenotype in the cells, and organoids from both conditions were analyzed for *LGR5, SMOC2* and *DKK4* by qRT-PCR. We observed a significant increase in the expression of *LGR5* (4.6-fold), *SMOC2* (78-fold), and *DKK4* (6992-fold) across the three adenomas (Fig. 7A). We further validated the protein expression of *LGR5* (Fig. 7B; Fig. S10) as well as the co-localization in adjacent histological sections of *LGR5* and *SMOC2* mRNA expression in colon organoids (Fig. 7C) and the co-localization of *LGR5* and *DKK4* in colon adenocarcinoma tissue (Fig. 7D) by *in situ* hybridization.

**Fig. 7.**
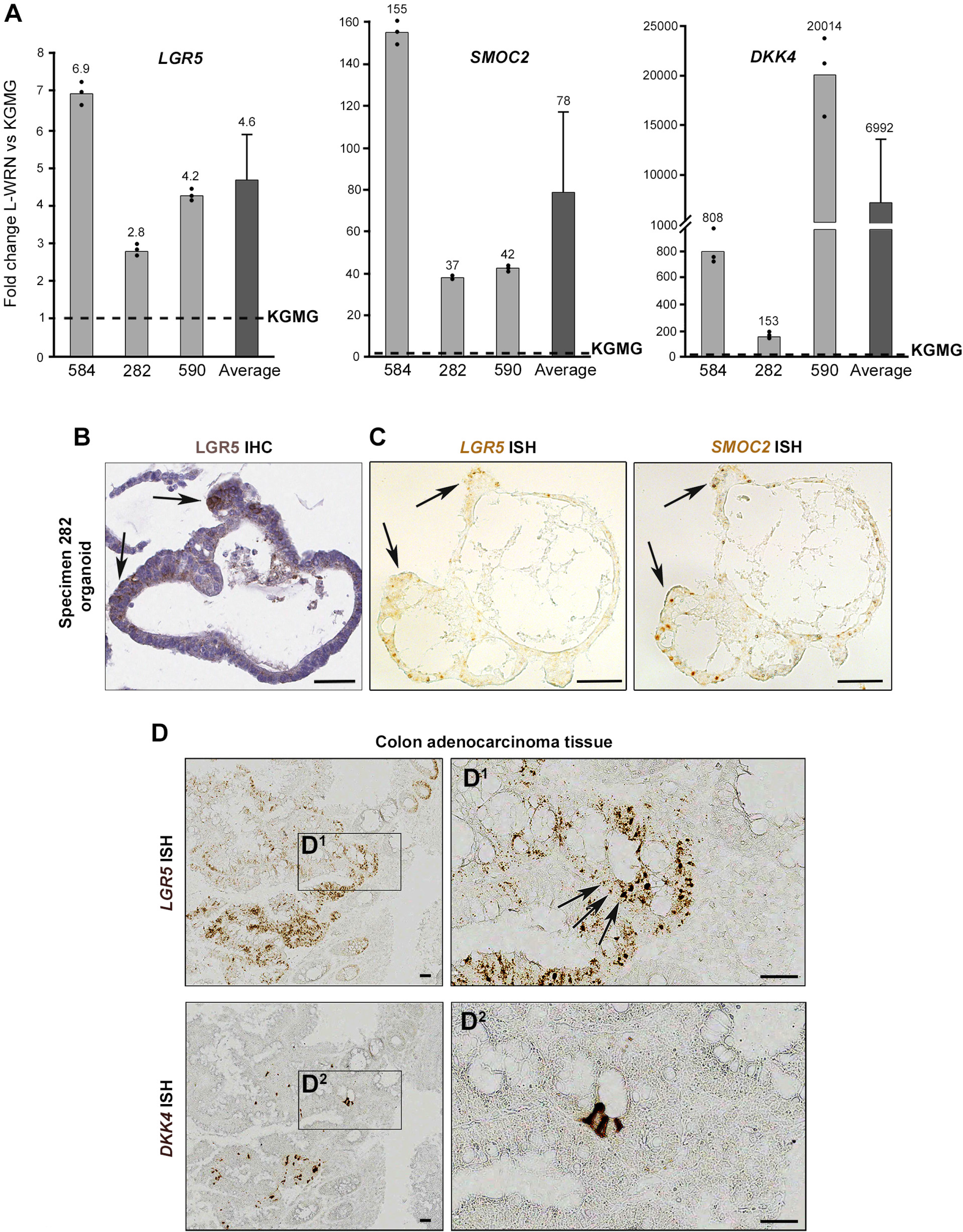
Association of *SMOC2* and *DKK4* expression patterns with Lgr5 expression. **(A)** Relative *LGR5*, *SMOC2*, and *DKK4* mRNA expression in L-WRN vs KGMG adenoma organoid cultures. Individual specimen qRT-PCR technical replicates; represents the mean±s.e.m. of n=3 biological replicates. **(B)** LGR5 immunohistochemistry staining and **(C)** *SMOC2* and *Lgr5* mRNA *in situ* hybridization (ISH) of formalin fixed paraffin-embedded (FFPE) adenoma 282 organoids cultured in L- WRN. Arrows designate localization of *Lgr5* and *SMOC2* in corresponding regions of consecutive serial sections. Scale bar, 50μM. **(D)** *DKK4* and *Lgr5* ISH images in consecutive FFPE cuts of a colon adenocarcinoma (specimen 815). Scale bar, 50μM.

Because we isolated enriched LGR5(+) cells from L-WRN-cultured adenoma organoids, we also wanted to isolate LGR5(+) cells directly from normal colon tissue to demonstrate robustness of the isolation and purification methods. Using the same dissociation methods as were employed for the adenoma organoid cultures, dissociated single cells from 3 independent normal cadaveric colon specimens were sorted for EpCAM (epithelial cell adhesion molecule) positivity and Lgr5 expression. LGR5(+) cells comprised 2.35%, 0.63%, and 0.76% of the EpCAM(+) cells in independent experiments. A representative distribution of LGR5(+) staining from a normal colon, demonstrates that fluorescent intensity for LGR5(+) events were 2-3 orders of magnitude above baseline (Fig. 8A). LGR5(+) cells were collected and subsequently analyzed by qRT-PCR for *LGR5* and the intestinal stem cell marker *OLFM4* (olfactomedin 4), which was recently shown to be expressed in normal human colon, and in adenoma (Jang et al., 2016). We identified an average of 7.5-fold increase in *LGR5* expression and 2.3-fold increase in *OLFM4* mRNA expression in the sorted LGR5(+) normal colon cells relative to the LGR5(−) cells (Fig. 8B). Further, protein expression of OLFM4 was confirmed by IF, which showed staining at the bottom 1/3^rd^ of the normal colon crypts (Fig. 8C).

**Fig. 8.**
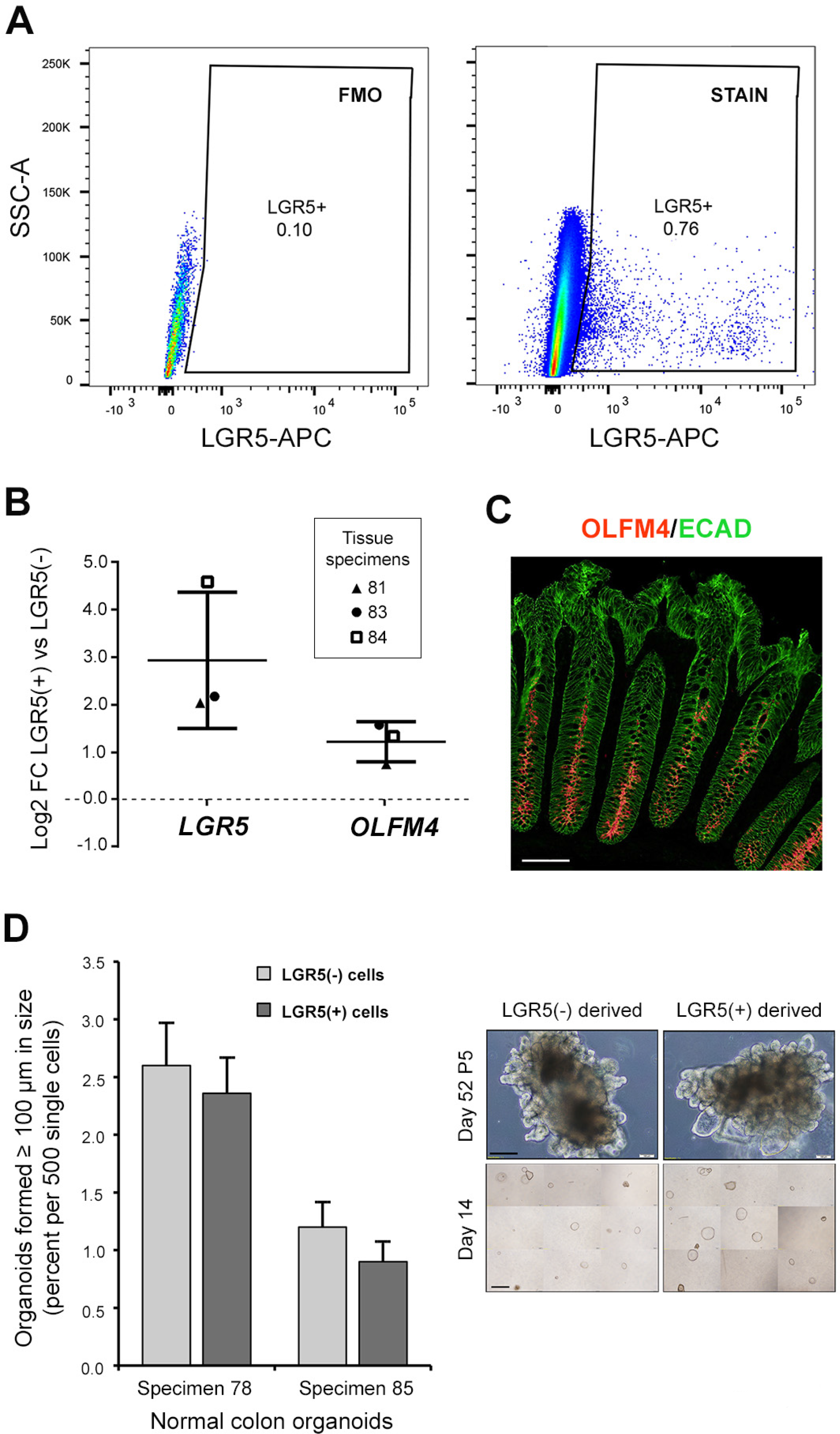
Identification, isolation and culture of patient-derived normal colon LGR5(+) cells. **(A)** Flow cytometry gating strategy for LGR5(+) cells isolated directly from warm autopsy normal colon tissue (specimen 84, a representative scatter plots of 3 patients; Table 1). Includes EpCAM(+) inclusion criteria, using a phycoerythrin (PE)-conjugated antibody (not shown). To set the LGR5-APC(+) gating, either magnet flow-through (specimen 81 & 83), as done with organoid sorts, or rigorous so-called fluorescence-minus-one controls [FMO; DAPI(−), EpCAM(+); specimen 84] were used. **(B)** mRNA expression by qRT-PCR of *Lgr5* and *OLFM4* across all 3 patient samples. Values represent mean ± SE. **(C)** OLFM4 immunofluorescent staining of normal colon crypts. Scale bar, 100μM. **(D)** Normal colon organoid cultures were established from warm autopsy resections. LGR5(+)/(−) cells were isolated from early passage organoids, sorted into culture medium and seeded into Matrigel to access organoid-forming efficiency. Mean organoid-forming efficiency from two normal colon organoid cultures (specimen 78, passage 5; and specimen 85, passage 3) sorted for Lgr5(−) vs. LGR5(+) cells (n=4 wells per fraction), error bars represent s.e.m. Representative images (right panels; specimen 85) of organoids formed from Lgr5(−) and LGR5(+) cells shown at day 14 and day 52 (passage 5). Scale bars, 200μM top panel; 1mm bottom panel.

Isolated LGR5(+) and LGR5(−) cells were sorted into medium and seeded into Matrigel to access organoid-forming efficiency (Fig. 8D). Those cells isolated directly from patient tissue did not initiate organoids under single cell culture conditions (n=3; data not shown). Instead, we established normal organoid cultures from the colon cadaveric resections (Table 1), and isolated LGR5(+) and LGR5(−) cells from early passage organoids. Organoids formed from sorted LGR5(+) and LGR5(−) cells and were passaged up to five times before terminating the experiment. Both LGR5(+) and LGR5(−) cells could form organoids with equal ability. The nearly equivalent capacity to form organoids between Lgr5(+) and LGR5(−) cells suggest that LGR5(−) cells, especially when cultured in a medium which drives WNT signaling (L-WRN), are capable of much plasticity, as other recent studies have recently elaborated (Richmond et al., 2015, Beyaz et al., 2016, Tetteh et al., 2016, Sasaki et al., 2016, Shimokawa et al., 2017, Yan et al., 2017, Jadhav et al., 2017).

## DISCUSSION

LGR5 is one of the few cell surface markers available for the identification and isolation of cycling stem cells in the colonic crypt. Efforts to generate effective antibodies to isolate these cells have largely been unsuccessful (Barker, 2014). Recently antibody drug conjugates (ADCs) have been developed to target cells expressing LGR5, demonstrating therapeutic potential (Gong et al., 2016, Junttila et al., 2015), but as part of these applications primary human cell isolation and characterization was limited (Hirsch and Ried, 2016). Thus, the role of Lgr5(+) stem cells in human intestinal development, homeostasis, and carcinogenesis remains poorly understood. The identification and isolation of human LGR5(+) adenoma cells, as well as the establishment of methods for the long-term culture and propagation of organoids enriched for these rare cells, as described in this study, provides some of the first insight into the characteristics of these cells in humans. Further, we confirm that our methods are robust for the identification and isolation of LGR5(+) cells from normal human colon tissues. Ideally, methods and data sets presented in this work will be widely utilized to overcome a major hurdle in the field of colorectal cancer stem cell biology.

Human primary intestinal organoid culture systems represent an unparalleled tool to study colorectal cancer biology (Matano et al., 2015) and to develop precision medicine platforms for targeted therapeutics (van de Wetering et al., 2015). Here, we report a living biobank of human normal colon, adenoma, and adenocarcinoma tissues. We used this biobank to study and isolate(+) cells from four genetically diverse adenoma-derived organoids. We found that LGR5 is expressed at higher levels in organoids in response to Wnt3a, Rspondin-3, and Noggin, and that this is associated with a cystic phenotype, consistent with other reports (Sato et al., 2011b, Farin et al., 2012, Matano et al., 2015, Drost et al., 2015, Onuma et al., 2013). While this report was in preparation, a study eloquently demonstrated that Mex3a marks a subset of LGR5(+) slow cycling intestinal cells (Barriga et al., 2017). Indeed, we show that *MEX3A* is among the upregulated genes in isolated human colon LGR5(+) cells. Interestingly, while gene expression associated with LGR5 expression in human adenoma significantly overlapped with *Lgr5*^HI^ stem cells in the mouse (Munoz et al., 2012), there were several examples where human expression patterns were opposite to that of the mouse. For example, *SEMA3B* and *PLCE1* are enriched in mouse *Lgr5*^HI^ cells, but were downregulated in human LGR5(+) adenoma cells (FDR=0.04 and 0.002 respectively). Determining whether differential expression of these genes reflects differences in human compared to mouse biology or adenoma compared to normal human colon biology represents an important future direction for this research.

Histological approaches that compared LGR5(+) cells from adenoma and adenocarcinoma samples with normal colon revealed a number of unexpected biological findings. While LGR5 expression significantly correlated with advanced stage in colorectal cancers, we surprisingly identified that high grade, poorly differentiated, tumors had lower LGR5 expression compared to lower grade tumors. This result corroborates a recent report of *LGR5* mRNA expression, quantified by *in situ* hybridization, in human colorectal cancers which also reported a decrease in *LGR5* expression in poorly differentiated tumors (Martin et al., 2017). We also identified the presence of stromal LGR5 staining, and confirmed the specificity of this signal with *Lgr5* mRNA ISH. We did not however identify obvious stromal staining in normal tissue. We identified that stromal staining of LGR5 was associated with cancer stage based on a cohort of independent samples on a tissue microarray. Others have identified the presence of multipotent LGR5(+) stromal cells in the oral mucosa and tongue of adult mice expressing the *LGR5*-EGFP reporter with a phenotype resembling neural crest cells (Boddupally et al., 2016). Further, Lee et al eloquently demonstrated recently through lineage tracing experiments in mice that LGR5(+) cells located in the lung alveolar mesenchyme support differentiation of the alveolar epithelium (Lee et al., 2017). Our finding is likely of high relevance in light of the identification of stromal gene expression having a strong correlation with colorectal cancer subtypes (Calon et al., 2015). Future work should focus on understanding the role of these putative LGR5 expressing stromal cells in colorectal carcinogenesis.

The ability to isolate LGR5(+) and LGR5(−) cells from human samples will enable future studies aimed at understanding human stem cells and cancer. In our functional studies, we found that single FACS-isolated LGR5(+) cells from normal colon organoids did not have increased organoid forming efficiency compared to LGR5(−) cells. These results are in line, however, with a recent report of the organoid forming efficiency of human LGR5(+) cells labeled genetically using Cas9/CRISPR, where the authors demonstrated a reemergence of *LGR5*-GFP(+) cells from the sorted *Lgr5*(−) fraction, and noted that organoid forming efficiency was not increased in the *LGR5*(+) cells when compared with *LGR5(*−) cells (Shimokawa et al., 2017), further highlighting the inherent plasticity of intestinal cells (Richmond et al., 2015, Beyaz et al., 2016, Tetteh et al., 2016, Sasaki et al., 2016, Yan et al., 2017, Jadhav et al., 2017). We anticipate that these varied approaches will help to move our understanding of human biology forward.

Transcriptional comparisons of LGR5(+) adenoma stem cells with published normal colonic RNA-seq data (Uhlen et al., 2015), identified genes strongly upregulated in LGR5(+) adenoma cells that were virtually unexpressed in the normal human colon, the highest being *DKK4*. DKK4 is a negative regulator of Wnt signaling that has been reported as both upregulated (Matsui et al., 2009, Pendas-Franco et al., 2008) and downregulated in CRC (Baehs et al., 2009). A recent study showed that DKK4 acts selectively to inhibit a subset of WNT ligands, but is proteolytically inactivated (Sima et al., 2016). Elevated *DKK4* expression is linked with CRC chemoresistance (Ebert et al., 2012) and metastasis (Chen et al., 2015), both processes associated with cancer stem cells. We identified that *DKK4* is significantly upregulated in colorectal cancers, compared to normal colon, in data from TCGA. The association with *LGR5* expression suggests that DKK4 could be a novel biomarker of colon adenoma stem cells or cells with high Wnt activity, and may prove useful for colon cancer prevention or treatment efforts.

Notably, our experiments used a two-step enrichment methodology to obtain purified cells (magnet followed by FACS), yet we still encountered a relatively small population of LGR5(+) cells. This could be explained, in part, by work showing rapid internalization of LGR5 from the plasma membrane to the trans-golgi network (Snyder et al., 2017). Thus, it is possible that our antibody-based methods will only recognize rare cells expressing the highest levels of LGR5 protein on the cell surface. On the one hand, this may preclude us from isolating cells that express LGR5 mRNA but that do not have abundant protein localized to the cell surface. On the other hand, this same caveat would enhance specificity for the sorted population of cells, as supported by our gene expression profiling.

Our study presents a blueprint for the identification, *in vitro* culture, isolation, and experimental characterization of human LGR5(+) cells. We anticipate our methods can be readily adapted for the isolation and characterization of the LGR5(+) cell populations in other human tissues including cochlea (McLean et al., 2017), hair (Jaks et al., 2008), kidney (Barker et al., 2012), liver (Huch et al., 2013), lung (Lee et al., 2017), mammary gland (De Visser et al., 2012), oral mucosa (Boddupally et al., 2016), prostate (Wang et al., 2015a), and stomach (Barker et al., 2010, Simon et al., 2012). Additionally, we expect that the organoid culture methods described here will provide an experimental platform for the development of novel chemopreventive and chemotherapeutic agents that target LGR5(+) stem cells.

## MATERIALS AND METHODS

### Establishment of organoid cultures from human colonic adenomas

Isolation of human colonic crypts and adenomas, and culture and maintenance of adenoma organoid cultures, were performed using our previously described protocol (Dame et al., 2014). Adenoma tissue was acquired by endoscopy and adenocarcinoma tissue was collected from colonic resections according to protocols approved by the University of Michigan Institutional Review Board (IRB; HUM00064405/0038437/00030020), with informed consent obtained from all subjects. Normal colonic tissues were collected from deceased donors through the Gift of Life, Michigan (HUM00105750). Deidentified human fetal intestinal tissue was obtained from the University of Washington Laboratory of Developmental Biology and approved by University of Michigan IRB (HUM00093465). Growth media used for organoid cultures included KGM-Gold™ medium (Lonza) (Dame et al., 2014) and L-WRN medium. KGMG is a serum-free epithelial medium containing a reduced calcium concentration of 0.15mM, hydrocortisone, epinephrine and pituitary extract, as well as some components common to L-WRN media, epidermal growth factor (although 1:1000 of L-WRN; 0.1ng/mL), insulin, and transferrin. L-WRN conditioned medium (Miyoshi and Stappenbeck, 2013) contains high levels of Wnt3a, R-spondin-3 and Noggin, with added 10mM Nicotinamide (Sigma-Aldrich). Adenoma organoid cultures were propagated long-term in KGMG. To drive organoids from a budding to cyst morphology, and to enrich for a “stemness” phenotype, cultures were switched from the reduced medium, KGMG, to the stem cell support medium, L-WRN, for 3-4 weeks. Consistent with previous reports, organoids formed cystic structures in the presence of Wnt ligand provided by L-WRN medium (Sato et al., 2011b, Farin et al., 2012, Matano et al., 2015, Drost et al., 2015, Onuma et al., 2013). Cultures were propagated in Matrigel (Corning) which was made to 8mg/mL in growth media, in 6-well tissue culture plates. Cultures were passaged every 4-7 days by digesting Matrigel in cold 2mM EDTA and plated on the first day with 10μM Y27632 (Miltenyi Biotec), a Rho-associated protein kinase (ROCK) inhibitor. For further organoid line details, including media components and concentrations, subculturing methods, genomic characterization, cell line authentication and mycoplasma monitoring, see supplementary Materials and Methods.

### Single Cell Isolation and Magnetic separation for LGR5(+) and LGR5(−) cells

Single cell suspensions of adenoma organoids or normal intact colon tissue were generated using the Tumor Dissociation Kit (human; Miltenyi Biotec) in combination with the gentleMACS Dissociator (Miltenyi Biotec) with protocol modifications. On the day of sorting, organoid cultures were treated with 10μM Y27632 for 2.5 hrs prior to harvest. The enzymes designated “H” and “R” were prepared in HBSS modified to 0.13mM calcium and 0.9mM magnesium (10% standard HBSS concentrations) to minimize differentiation of the epithelial cells while supporting enzymatic activity. All plasticware, including cell strainers and columns, were 0.1% BSA-coated, and buffers contained 5-10μM Y27632. Cells were labeled with anti-LGR5 MicroBeads (human; Miltenyi Biotec) and the LGR5(+) cells were enriched by Magnetic-Activated Cell Sorting (MACS). Cells were applied through a cold BSA-coated 20μM cell strainer (CellTrics) to LS Columns (Miltenyi Biotec) in the above HBSS buffer containing 200 Kunitz units/mL DNAse (Sigma-Aldrich), 0.5% BSA in DPBS (Sigma-Aldrich), and 5μm Y27632. The magnet-bound positive and flow-through negative fractions were analyzed and isolated by FACS. For further details, regarding single cell preparation and MACS, see supplementary Materials and Methods.

### Flow Cytometry

Cells were analyzed on a LSRII cytometer (BD Biosciences) and sorted on a MoFlo Astrios (Beckman Coulter). Events first passed through a routine light-scatter and doublet discrimination gate, followed by exclusion of dead cells using 4’,6-diamidino-2-phenylindole (1μM DAPI dilactate; Molecular Probes). Gating strategy set the APC-positive cell population at 0.05-0.1% of the viable MACS flowthrough cells, LGR5(FT+) (Fig. 3C bottom panels; all FACS RNA sequencing-derived data; Fig. 8A, specimen 81, 83). Gating strategy set the APC-positive cell population at 0.05-0.1% of the viable FMO- APC check reagent control for the analysis of samples that were not MACS processed (Fig. 3C top and middle rows; Fig. S2A). Flow cytometric data analysis was performed using Winlist 3D software (Verity) and FlowJo vX.0.7 (Tree Star). See supplementary Materials and Methods for further details.

### High Throughput RNA Sequencing (RNA-seq)

RNA was extracted from sorted cells using the RNeasy Micro Kit (Qiagen) with on-column DNase digestion. Due to the small number of cells following FACS and corresponding low level of input RNA, we depleted ribosomal RNAs with RiboGone (Takara Clontech) and prepared sequencing libraries utilizing the SMARTer Stranded RNA-Seq kit (Takara Clontech,). Libraries were multiplexed over 2 lanes and sequenced on a HiSeq sequencer (Illumina). For further details, including alignment and quality control, differential expression testing, and KEGG pathway analyses, see supplementary Materials and Methods.

### Comparison to previously published datasets

RNA-seq data from normal colon (Tissue Based Map of the Human Proteome Science REF) and differential gene expression between the LGR5(+) cells and normal colon were calculated as described above (Fig. 6B, C). Overlap between genes overexpressed in LGR5(+) cells with the previously reported *Lgr5*+ ISC gene signature (Muñoz et al., 2012) was calculated by identifying the overlap between genes present in both the Munoz et al ISC gene expression signature and genes identified as upregulated in LGR5(+) adenoma cells (Fig. 6A; Table 2; Table S2). Statistical significance of the overlap between these gene signatures was calculated using the hypergeometric distribution.

### LGR5 Immunohistochemistry

Formalin fixed, paraffin sections were cut at 5-6 microns and rehydrated to water. Heat induced epitope retrieval was performed with FLEX TRS High pH Retrieval buffer (pH 9.01; Agilent Technologies, 154 #K8004; Santa Clara, CA) for 20 minutes (Fig. 1A, C and Fig. 3A). After peroxidase blocking, the antibody LGR5 rabbit monoclonal clone STE-1-89-11.5 (Miltenyi Biotec) was applied at a dilution of 1:50 (Fig. 1A, C) or 1:100 (Fig. 3A) at room temperature for 60 minutes. The FLEX HRP EnVision System (Agilent Technologies) was used for detection with a 10 minute DAB chromagen application. Note, sections freshly cut were compared to those that were stored at room temperature for 4 weeks, and showed more robust LGR5 staining (data not shown). The colon cancer tissue microarray (Fig. 3A; 2 normal, 3 adenoma, 70 adenocarcinoma; 2 core samples per specimen) was freshly cut and provided by BioChain Institute, Inc.

### LGR5 Immunohistochemistry scoring for staining intensity in the epithelium and in the stroma

The TMA, along with five additional FFPE normal colon samples from warm cadaveric colon resections, were scored for staining intensity in both the epithelium and then separately in the stroma (Fig. 3). Scoring was conducted by two independent viewers on blinded samples at 8X and 20X magnification. Scoring key: 0 = non-specific or < 1%; 1 = 1-10% or only evident at 20X magnification; 2 = 10-50% or light diffuse staining >50%; 3 = >50%. Stage I (n=24), Stage II (n=24), and Stage III (n=21) tumors were compared to normal colon (n=7) and adenoma (n=3). TMA cancers with grades I & I-II were grouped, termed “Grade I”, and cancers grade II-III & III were grouped, termed “Grade III” for further analyses. LGR5 stromal and epithelial staining for adenomas (n=3), cancer Grade I (n=20), II (n=38), and III (n=10) were compared to normal colon tissue (n=7). For additional details regarding IHC staining and scoring see supplementary Materials and Methods.

### LGR5 Immunofluorescence

Rehydrated sections were retrieved with R-Buffer B (pH 8.5; Electron Microscopy Sciences) in a pressurized Retriever 2100 (Electron Microscopy Sciences) overnight. Slides were blocked for 1 hour with 0.5% Triton X-100 and 5% donkey serum and were incubated overnight at 4°C with the following antibodies in 0.05% Tween 20 and 5% donkey serum: anti-LGR5 clone 89.11 (Fig. 1C^3^C^4^); anti-OLFM4 antibody (Abcam) (Fig. 8C); anti-defensin5a (Abcam) (Fig. S1A); anti-E-cadherin (BD Bioscience) (Fig. 8C); and anti-E-cadherin (R&D) (Fig. S1A), followed by 1 hour room temperature incubation with the following secondaries: LGR5 plus DAPI (Fig. 1C^3^/C^4^; Fig. 2A) and OLFM4 (Fig. 8C) secondaries (biotinylated secondary donkey anti-rabbit; Jackson Immuno Research Labs); defensin5a (Fig. S1A) and E-cadherin secondary (Fig. 8C) (donkey anti-mouse 488; Jackson Immuno Research Labs); E-cadherin secondary (Fig. S1A) (donkey anti-goat 647; Jackson Immuno Research Labs). LGR5 and OLFM4 were amplified with the SK4105 TSA Kit with Alexa 594 tyramide (Invitrogen). For additional IF staining details see supplementary Materials and Methods.

### Further Specificity Studies of LGR5 Antibodies and Development of Procedures for Enrichment of LGR5(+) cells by Magnetic-activated Cell Sorting (MACS)

Western analysis was performed with 1881 LGR5(+), wild type 1881 LGR5(−), specimen 282 and 14881. After protein transfer from a Tris-Glycine 4-20% gradient gel, the membrane was incubated with primary antibody, 0.1μg/mL rabbit anti human LGR5 STE-1-89-11.5 (Miltenyi Biotec), in 10mL TBS/Tween/milk powder overnight at 4°C, followed by incubation with secondary HRP goat anti rabbit IgG (Cell Signaling Technology). Flow cytometry spike-in experiments (Fig. S2B) were performed with 1881 LGR5(+) and 1881 LGR5(−) by mixing at varying proportions and analyzing on a LSRII cytometer (BD Biosciences) before and after magnetic separation with rat monoclonal antibody anti-human LGR5 clone 22H2.8 magnetic bead-conjugate (Miltenyi Biotec), with the allophycocyanin (APC) anti-bead check reagent to recognize the bound magnetic beads (Miltenyi Biotec). Detailed information regarding the culture and transfection of the 1881 cell lines, validation of 1881 *LGR5* transfection stability by qRT-PCR, flow cytometry/MACS methods, and western analysis, see supplementary Materials and Methods.

### Transduction of organoids with lentiviral TCF/LEF GFP reporter and ImageStream Analysis

Colon adenoma specimen 282 organoids were transduced with a lentiviral TCF/LEF GFP reporter (Qiagen) based on Koo et al (Koo et al., 2013) with the major modifications being that 1) organoids were mechanically dissociated in the absence of enzymatic digestions (to avoid single cells), and 2) that incubations were done at 4°C. Organoids were passaged under puromycin selection and transitioned to cystic morphology with L-WRN medium. Single cells were analyzed and imaged with the Amnis ImageStreamX Mark II (EMD Millipore) for co-expression of *TCF/LEF*-GFP and LGR5-APC fluorescence (Fig 5B,C). For further transduction and ImageStream details see supplemental Materials and Methods.

### Association of SMOC2 and DKK4 expression with LGR5 expression

*LGR5*, *SMOC2*, and *DKK4* mRNA expression were assessed by qRT-PCR and presented as increased expression in L-WRN relative to KGMG, with three adenomas organoids 282, 584, and 590, after transitioning from KGMG to L-WRN. *In situ* hybridization (ISH) was preformed to assess of localization of *SMOC2* and *DKK4* relative to *LGR5*. ISH was performed using the RNAscope 2.0 HD detection kit (Advanced Cell Diagnostics, ADC) according to the standard provided protocol (Fig. 1B, D; Fig 7C, D), and all probes were designed by ADC. See supplemental Material and Methods for further qRT-PCR and ISH details including primer/probe sequences.

### Normal Colinic Crypt LGR5(+) Cell Isolation and Culture

Crypts were isolated from three warm autopsy normal colon specimens (Table 1) and processed as previously described (Dame et al., 2014), a modification of Sato et al (Sato et al., 2011a), and normal colon organoid cultures were established (Table 1). LGR5(+)/(−) cells were isolated from early passage organoids (P3 and P5) and sorted (without MACS) into growth medium containing 0.25mg/mL Matrigel. The flow cytometry strategy was as above, with the added gating/analysis for EpCAM expression using a phycoerythrin (PE)-conjugated antibody (BioLegend). The medium composition was roughly based on previous mouse small intestine/colon single cell organoid-forming efficiency studies (Yin et al., 2014, von Furstenberg et al., 2014, Wang et al., 2013, Shimokawa et al., 2017, Jung et al., 2015). Sorted cells were seeded into 4 wells at 500 cells/28μL Matrigel/well (24-well plate). At day 14-17, images were taken and organoids over 100μm were counted to access organoid-forming efficiency (Fig. 8D). Cultures were passaged for approximately 2 months with cryopreservation before ending. For further details including FACS of normal cells and single cell medium components, see supplemental Material and Methods.

### Statistical Analyses

For IHC analyses, LGR5 stromal or epithelial staining intensity categories were plotted by tumor stage and grade using boxplots. Differences in LGR5 staining in the stroma or epithelium by stage or grade were quantified by linear regression, treating epithelial or stroma staining intensity (0, 1, 2, or 3) as a continuous dependent variable and either tumor stage or grade as a categorical independent variable, setting normal tissue as the reference group. For qRT-PCR analyses, differences between biological replicates across experimental conditions were assessed using t-test. For both IHC and qRT-PCR analyses, differences were considered statistically significant at *p* < 0.05. Statistical methodology for genomic variant characterization and RNA-sequencing differential expression analyses can be found in the Supplemental Information.

**Data Access:** The RNA-seq and genomic variant raw data are publically available at ArrayExpress under accession number E-MTAB-4698.

## Acknowledgments

We thank Kelly D. Maynard, Veda Yadagiri, Bodrul Islam, Kevin Kim, and Jessica Zhang for their support of the GISPORE Organoid Core facility; Missy Tuck, Brian Kleiner and Kim Gonzalez, clinical coordinators, the University of Michigan Hospitals, for their assistance in obtaining adenoma biopsies; Kristina Fields and Alan Burgess of the Research Histology and IHC Core; Dave Adams and Michael Dellheim of the UM BRCF Flow Cytometry Core; Deborah Postiff and Jackline Barikdar of the Tissue Procurement Core; Jeanne Geskes, Robert Lyons, and Melissa Coon, University of Michigan DNA Sequencing Core Facility; Ashwini Bhasi of the University of Michigan Bioinformatics Core; Robin Kunkel for graphic design, Department of Pathology; and Kaycee White and Muhammad N Aslam of the Histomorphometry Core, Department of Pathology, for their ScanScope service.

## Author contributions

Conceptualization: MD, DA, SM, JC, JS, YS, XX, JS, BS; Methodology: MD, DA, SM, OH, PO, DAg, AB; Investigation: MD, DA, SM, PD, PO, AC, XX, JL, EK, GN, DH, AM, YT, DAg, CA; Analysis: JC, JS, HA, JL, PO; Resources: LS, OH, JS, CA; Original Draft: JC, MD, DA; Review-Editing: JS, YS, DB, LS, MW, JV; Supervision JC, JS, YS, DB, MW, JV. All authors had access to the study data and reviewed and approved the final manuscript.

## SUPPLEMENTAL EXPERIMENTAL PROCEDURES AND FIGURES

### Establishment of Organoid Cultures from Human Colonic Adenomas

The isolation and culture procedure was based on recently described methodology for establishing and maintaining unlimited growth of human colonic adenoma tissue in organoid culture (Dame et al., 2014) founded in the original work of Sato *et al* (Sato et al., 2011a). Use of this procedure has generated a working cryorepository of adenoma organoids (Table 1). Tissue was collected from subjects recruited through the Molecular Pathology and Biosample Core (MPBC) as part of the GI SPORE at the University of Michigan. Colonic tissue was acquired by endoscopy (adenomas) or from colonic resections (adenocarcinomas) and according to protocols approved by the University of Michigan Institutional Review Board (IRB; HUM00064405/0038437/00030020), with informed consent obtained from all subjects. Adult small intestinal and colonic tissue was collected from deceased donors through the Gift of Life, Michigan (University of Michigan IRB HUM0010148810). Fetal small intestinal tissue was obtained from the University of Washington Laboratory of Developmental Biology and approved by University of Michigan under IRB approval (HUM00093465).

Two growth media were used. The patient-derived adenoma organoid specimens selected for this study were established and maintained in continuous culture in a reduced medium, KGM-Gold™, (#195769, Lonza, Walkersville, MD; KGMG) (Dame et al., 2014), a serum-free epithelial medium containing a reduced calcium concentration of 0.15mM (with CaCl_2_; PromoCell GmbH; #34006; Heidelberg, Germany), hydrocortisone, epinephrine and pituitary extract, as well as some components common to L-WRN media (below), epidermal growth factor (although reduced, 0.1ng/mL), insulin, and transferrin; and including the antimicrobials 25μg/mL gentamicin (Gibco, #15750060; Grand Island, NY) and 50μg/mL Primocin (InvivoGen, #ant-pm-1; San Diego, CA). The second medium, L-WRN medium, was employed to drive the budding organoids into a cystic morphology, high in Wnt3a, R-spondin-3 and Noggin, all provided by a conditioned medium from stably transfected support L-cells as described by Miyoshi (Miyoshi and Stappenbeck, 2013). The complete L-WRN medium contained Advanced DMEM/F-12 (Gibco, 12634028), 2mM GlutaMax (Gibco, #35050-061), 10mM HEPES (Gibco, #15630080), N-2 media supplement (Gibco; #17502048), B-27 supplement minus vitamin A (Gibco; 12587010), 50 units/mL penicillin, 0.05 mg/mL streptomycin (Gibco, #15070063), 50μg/ml Primocin (InvivoGen; San Diego, CA), 1mM N-Acetyl-L-cysteine (Sigma-Aldrich, A9165; St. Louis, MO), and 100ng EGF/mL (R&D Systems, Inc., 236-EG; Minneapolis, MN), as well as the addition of 10mM Nicotinamide (Sigma-Aldrich, N0636). To drive the cultures to a cyst morphology, cultures were gradually switched from KGMG to L-WRN for three to four weeks (3 days KGMG post-passage, 5 days 50/50 L-WRN; 5 days 75/25; 100% L-WRN until predominately cystic). Consistent with previous reports, organoids formed cystic structures in the presence of Wnt ligand provided by L-WRN medium (Onuma et al., 2013, Drost et al., 2015, Sato et al., 2011b, Farin et al., 2012, Matano et al., 2015).

Growth of normal colon organoids and some adenoma specimens required the above L-WRN medium with additional growth factors: 10μM SB202190 (Sigma-Aldrich; S7067) and 500nM A83-01 (R&D Tocris, #2939) (Sato et al., 2011a). Further, a subset of these adenoma showed inhibition of growth with the p38 MAP kinase inhibitor, SB202190 (Sigma-Aldrich; #S7067), as described by Fujii et al (Fujii et al., 2016). 10mM Nicotinamide was used at the initial establishment of the specimens in organoid culture, and then removed for continued expansion (Bartfeld et al., 2015).

Cultures were propagated in Matrigel (Corning, #354234; Bedford, MA) which was diluted to 8mg/mL in growth media and grown on 6-well tissue culture plates (USA Scientific CytoOne, #CC7682-7506; Ocala, FL). All plasticware, excluding culture plates, were coated in 0.1% bovine serum albumen in DPBS (Sigma-Aldrich; #A8806) to prevent tissue/organoids/cells from adhering to surfaces. Cultures were passaged every 4-7 days by digesting Matrigel in cold 2mM EDTA in DPBS and plated the first day with the Rho-associated protein kinase (*ROCK*) inhibitor, 10μM Y27632 (Miltenyi Biotec GmbH, #130-104-169; Bergisch Gladbach, Germany). For normal colon and select adenoma grown in L-WRN, 2.5μM CHIR99021 (Miltenyi Biotec, #130-103-926) was also added at passaging.

### Genomic Characterization of Organoid Cryorepository Specimens

DNA was extracted from Allprotect Tissue Reagent (Qiagen, #76405; Germantown, MD) preserved samples or directly from fresh organoid preparations with the All prep DNA/RNA/protein mini kit (Qiagen, #80004). For specimens 574 and 584, DNA was isolated from FFPE source tumor with the QIAamp DNA FFPE Tissue Kit, (Qiagen, #56404).

The QIAseq Targeted DNA colorectal cancer panel (Qiagen, #DHS-002Z) was used to determine the presence of somatic and germline variants (Table 1). The panel is compatible with Illumina sequencing technology and covers 71 different oncogenes and tumor suppressor genes often mutated in colorectal cancers. The assay was performed using 80ng of purified gDNA following the vendor’s manual. The libraries were sequenced on a HiSEQ 2500 using V2 chemistry (Illumina Inc.; San Diego, CA). The sequencing data was demultiplexed and the Fastq files generated using Casava. The fastq files were imported into the QIAseq DNA enrichment portal for alignment and variant analysis. The targets were covered at an average read depth of 2000x with a coverage uniformity at 20% of the mean over 99%. Somatic variants were manually inspected in the integrative genomics viewer (IGV, Broad Institute). The variants of low quality or with an allele frequency below 5% were filtered out.

Whole exome sequencing was performed for specimens 590 and 14881 (Table 1). Briefly, DNA was fragmented to 250bp using standard Covaris sonication. Fragmented DNA was prepared as a standard Illumina gDNAlibrary using IntegenX reagents. Then the samples were PCR amplified, libraries were checked for quality and quantity. Samples then underwent exome capture (using the Roche Nimblegen SeqCap EZ v3.0 according to the manufacturer’s protocols (Roche Nimblegen, Indianapolis, IN). The SeqCap EZ Human Exome Library v3.0 covers 64 Mb of coding exons and miRNAs. Captured libraries were sequenced on a HiSeq 2500 using 100 Cycle paired end reads. The exome sequencing data was analyzed by the variant calling pipeline developed by the University of Michigan Bioinformatics Core. For each of the samples, paired-end reads were aligned to the hg19 reference genome using BWA v0.7.8 (Li and Durbin, 2009), followed by removal of sequence duplicates using PicardTools v1.79 (http://picard.sourceforge.net), local realignment around INDELs and base quality score recalibration using GATK v3.2-2 (DePristo et al., 2011). Read coverage on exome capture target regions was calculated using BEDTools v2.20.1 (Quinlan and Hall, 2010). Normal-Tumor paired alignment files were submitted to MuTect v1.1.4 (Cibulskis et al., 2013), Strelka v1.0.14 (Saunders et al., 2012), and Varscan v2.3.7 (with its false-positive filter) (Koboldt et al., 2012) for the detection of somatic and germline SNPs and INDELs. Candidate variant calls across all samples and patients were merged using Jacquard (Gates et al., 2015) into a single VCF file that included all variant loci whose filter field passed in MuTect or Strelka or VarScan (VarScan calls were limited to somatic variants confirmed in false-positive filter). Variants were annotated using SnpEff v4.0/hg19 (Cingolani et al., 2012), dbNSFP v2.4 (Liu et al., 2011, Liu et al., 2013), dbSNP v138, and 1000 Genomes v3 (Abecasis et al., 2012). For variants associated with multiple effects or multiple transcripts, a single “top effect” annotation was nominated based on annotation confidence, predicted impact, gene region, and transcript length. Common variants (at or above 5% overall population allele frequency as reported by 1000 Genomes) were excluded. Subsequent variant calling was as follows: rejected Impact Ranks low & modifier and accepted high & moderate; rejected Population Category common; excluded dbNSFP when damaging was equal to zero for all nine data bases, and selected allelic frequency ≥ 0.15.

### Organoid line authentication and mycoplasma monitoring

Organoid lines were authenticated by short tandem repeat (STR) analysis at the University of Michigan DNA Sequencing Core with the AMPFLSTR Identifier Plus Assay (Applied Biosystems) to identify human genomic DNA for 15 tetranucleotide repeat loci and the amelogenin gender determination marker run on the 3730XL Genetic Analyzer (Applied Biosystems). Adenoma and adenocarcinoma lines were also matched by variant profiling (above) to their source tissue, showing a high degree of concordance up to 9 months in extended culture. Cultures are periodically monitored for contamination by enzymatic activity with the Lonza MycoAlert Kit as a service of the UMICH Transgenic Animal Model Core.

### Further Specificity Studies of LGR5 Antibodies and Development of Procedures for Enrichment of LGR5(+) cells by Magnetic-activated Cell Sorting (MACS)

Mouse pre-B cell lymphoma 1881 cells transfected with his-tagged *LGR5* [1881(+)] or wild type [1881(−)] (provided by Miltenyi Biotec) were cultured in suspension in RPMI (Gibco, #21870-076) containing 20mM HEPES, 2mM L-glutamine, 10% FBS (Hyclone characterized, GE Healthcare Life Sciences, #SH30071.03; Marlborough, MA), 50 units/mL penicillin, 0.05 mg/mL streptomycin, 1mM sodium pyruvate (Gibco, #11360070), and 50μM β-mercaptoethanol (Gibco, #21985023). Transfected cells were selected using 15μg/ml Blasticidin S (Gibco, #R210-01).

#### *LGR5* quantitative real-time PCR of 1881 LGR5(+) and 1881 LGR5(−) cells

To confirm transfection stability, the 1881 LGR5(+) and 1881 LGR5(−) cells were assayed for *LGR5*, and visualized with agarose gel analysis of qRT-PCR amplified products. mRNA expression of 1881 LGR5(+) and 1881 LGR5(−) cells was quantified by qRT-PCR (Fig. S1B; three biological replicates). Total RNA was extracted using TRIzol Reagent (Invitrogen, #15596-026; Carlsbad, CA) according to the manufacturer’s recommendations. Complementary DNA was generated using TaqMan Reverse Transcription Reagents (Applied Biosystems, N808024; Carlsbad, CA). Quantitative real-time PCR was performed using Brilliant II SYBR Green Master Mix (Agilent technologies, #600828; Santa Clara, CA) on a StepOnePlus Real-Time PCR System (Applied Biosystems). All reactions were run with 40 cycles of 95°C for 10 minutes, 94°C for 30 seconds, 60°C for 30 seconds, and 72°C for 60 seconds, followed by a melt curve of 95°C for 10 seconds, 60°C for 1 minute and then increasing at 0.5 degree increments to 95°C. Gene expression analysis was normalized to the housekeeping gene *ACTB* and values were plotted as arbitrary units. The qRT-PCR amplified products (Fig. S1B) were analyzed by running on a 1.5% Agarose gel (Bio-Rad Laboratories, #1620137; Hercules, California) to confirm the target amplicons size and to indicate that there were no non-specific amplification products. Primer sequences were as follows: *hLGR5* (Finkbeiner and Spence, 2013): CAGCGTCTTCACCTCCTACC (Forward primer); TGGGAATGTATGTCAGAGCG (Reverse primer); 126 bp band size. *mACTB* (Riera et al., 2014): CTAAGGCCAACCGTGAAAAG (Forward primer); CCAGAGGCATACAGGGACA (Reverse primer); 104 bp band size.

**Fig. S1 (to Fig. 1).**
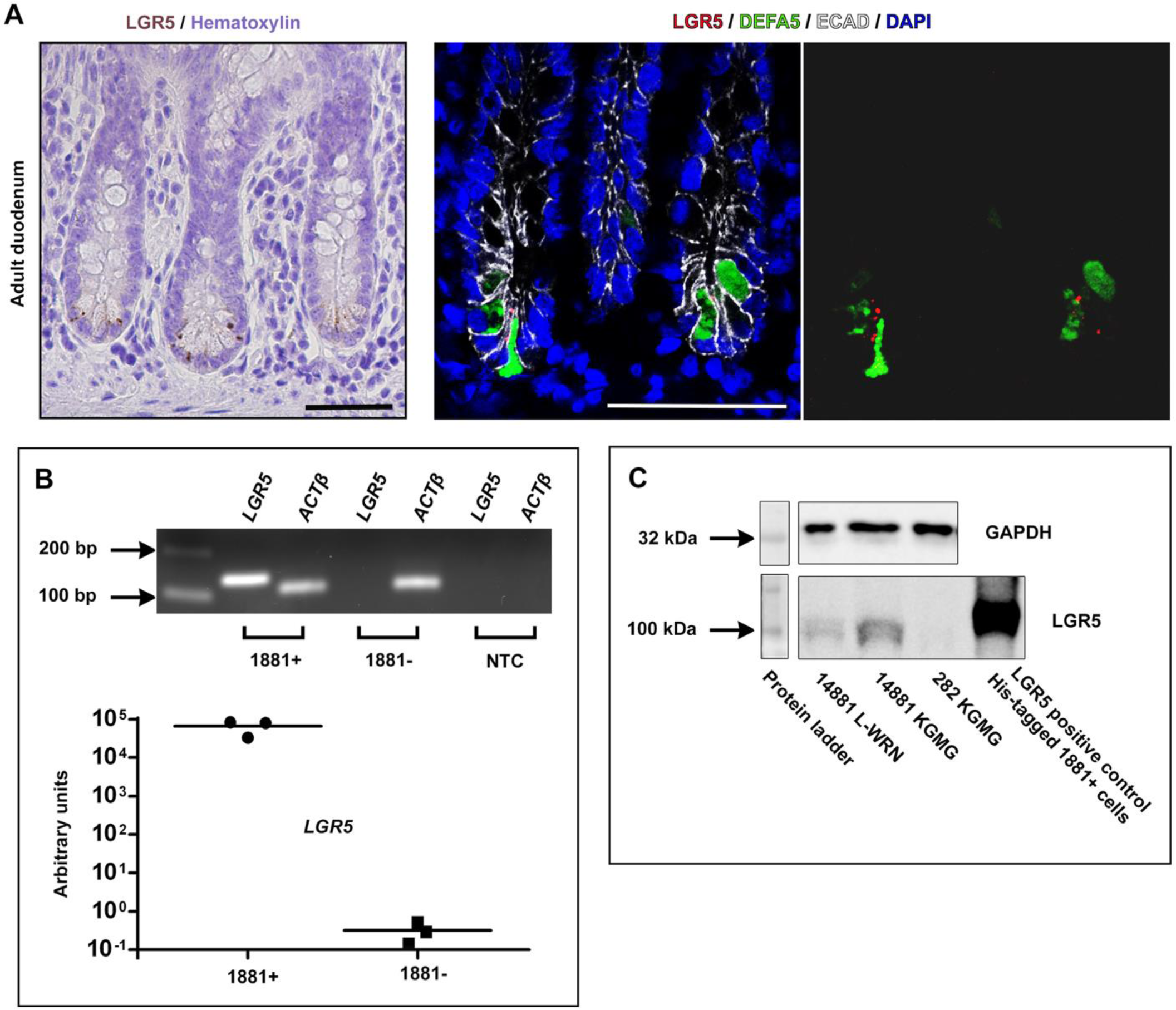
Further specificity analysis of human LGR5 antibody. **(A)** LGR5 immunohistochemical and immunofluorescent staining in adult duodenum showed weak punctate LGR5 expression, while adjacent cells stained with the paneth cell marker DefensinA5 (DEFA5) (middle and right panel). Scale bars, 50μM. **(B)** Mouse 1881 lymphoma cells were previously transfected with human *Lgr5* [1881(+); Miltenyi Biotec]. To confirm transfection stability, the 1881 LGR5(+) and 1881 Lgr5(−) cells were assayed for *Lgr5* expression (n=3 biological replicates), and visualized with agarose gel analysis of qRT-PCR amplified products. Line represents mean values. (C) Measurement of human LGR5 protein expression by Western blotting with the rabbit monoclonal anti-human LGR5 antibody clone STE- 1-89-11.5 in 1881(+) cells, 282 adenoma organoids cultured in KGMG, and 14881 adenoma organoids cultured in KGMG or L-WRN.

#### Flow cytometry spike-in experiments

Cells were analyzed on a LSRII cytometer (BD Biosciences) to confirm LGR5 antibody specificity and validate the magnetic bead enrichment strategy (magnetic-activated cell sorting, MACS) (Fig. S2B). 1881 LGR5(+) and wild type 1881 LGR5(−) cells were mixed at varying proportions and analyzed by flow cytometry before and after magnetic separation with rat monoclonal antibody anti-human LGR5 clone 22H2.8 conjugated to magnetic beads (Miltenyi Biotec, #130-104-072), with the allophycocyanin (APC) anti-bead check reagent conjugate to recognize the bound magnetic beads (Miltenyi Biotec, #130-098-892). Detailed flow cytometry/MACS methods are as described in the below methods section, “Single Cell Isolation and Magnetic separation for LGR5(+) and Lgr5(−) cells”.

**Fig. S2 (related to Fig. 3).**
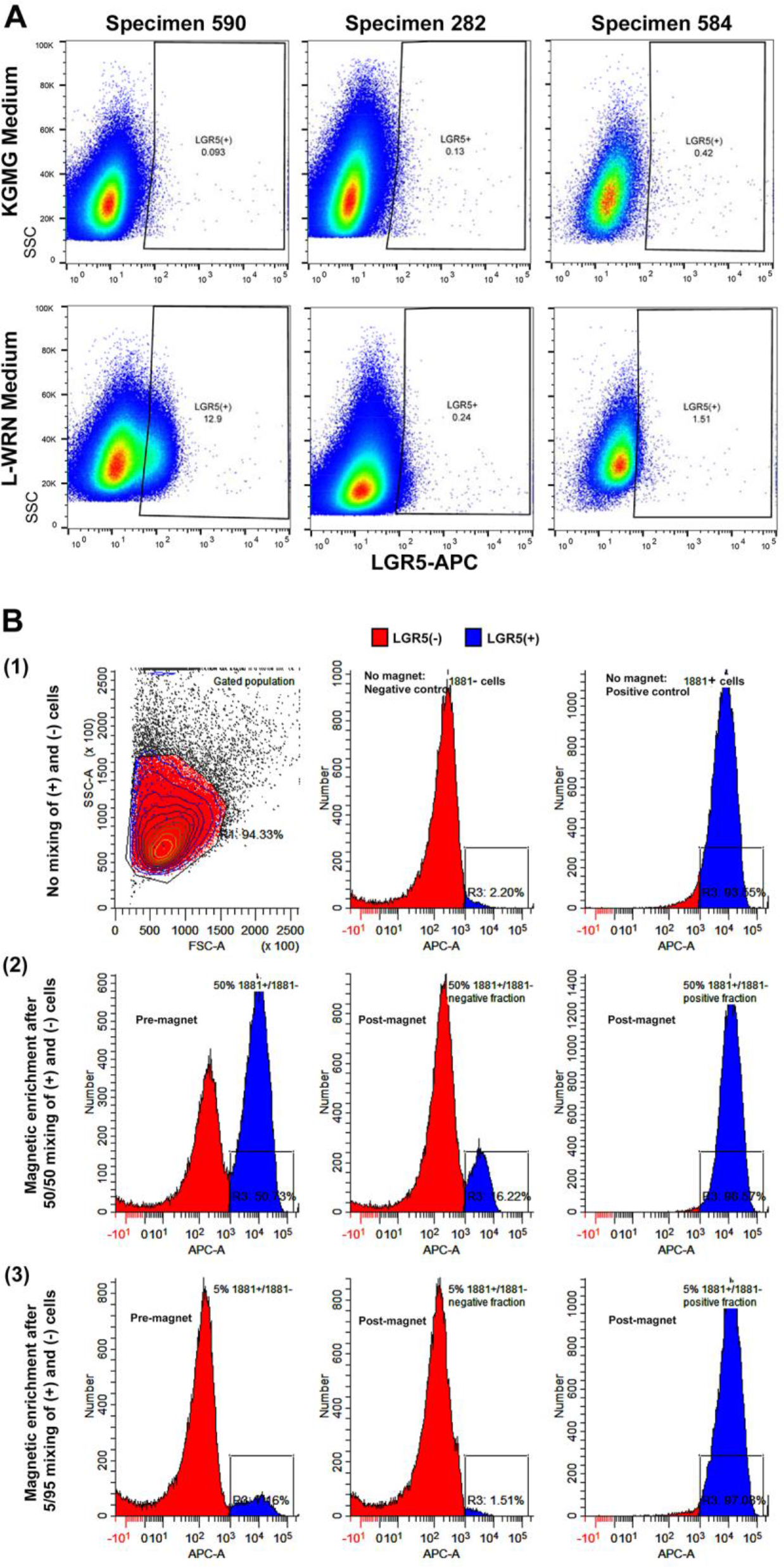
Development of procedures for enrichment of LGR5(+) cells; L-WRN culture and magnetic-activated cell sorting (MACS). **(A)** Flow cytometric analysis for enhanced number and fluorescent intensity of LGR5(+) cells resulting from organoids cultured in the reduced medium KGMG and then transferred for 34 weeks to L-WRN medium. **(B)** Flow cytometry spike-in experiments to confirm LGR5 antibody specificity and to validate the magnetic bead enrichment strategy. (1) Pure populations of 1881 LGR5(+) and 1881 Lgr5(−) cells were analyzed by flow to establish gating strategy. 1881 LGR5(+) and 1881 Lgr5(−) cells were mixed in the proportions, (2) 50/50 and (3) 5/95; and analyzed by flow cytometry before (left column) and after magnetic separation (middle/right columns) with rat monoclonal anti-human LGR5 antibody clone 22H2.8.

### Western analysis

Whole-cell extracts were isolated, separated and transferred to nitrocellulose as previously described (Xue et al.). Cell lysates were prepared with the following: 150mM NaCl, 50mM Tris/ HCl pH8, 1% NP-40, 0.5% Triton-X 100, Protease inhibitor complete (Roche Life Science, # 04 693 116 001; Penzberg, Germany), fresh 2μg/mL aprotinin, fresh 10μg/mL leupeptin, fresh 100μg/mL PMSF, at 200μl buffer-mix per 1x10^7^ cells and incubated for 30 minutes on ice. Samples were centrifuged at 13,000xg for 15 minutes at 4°C and supernatant transferred to a fresh tube to determine the protein concentration (Bradford-assay). Before loading, samples were boiled for 5 minutes at 95°C. 15p,g of sample protein was loaded to a Tris-Glycine 4-20% gradient gel (using Tris Glycine SDS running buffer). Protein was transferred to a nitrocellulose membrane and the membrane was blocked in TBS/Tween/milk powder (150mM NaCl; 50mM Tris pH 7,0; 0.1% Tween 20; 5% skim milk powder) for 1 hour at room temperature. The membrane was incubated with primary antibody 0.1μg/mL rabbit anti human LGR5 STE-1-89-11.5 (Miltenyi Biotec, #130-104-945) in 10mL TBS/Tween/milk powder overnight (shaking slightly) at 4°C, followed by 3X washes for 5 minutes with TBS/Tween (150mM NaCl; 50mM Tris pH 7,0; 0.1% Tween 20) (shaking slightly). The membrane was incubated with secondary antibody (HRP goat anti rabbit IgG, 1:2000; Cell Signaling Technology; Boston, MA), diluted in 10mL TBS/Tween/milk powder for 1 hour at room temperature (shaking slightly), followed by 3X washes for 5 minutes with TBS/Tween, 1X wash with deionized water for 5 minutes. The chemiluminescent signal was detected using Immobilon™ Western Kit (EMD Millipore) (Fig. S1C).

### LGR5 Immunohistochemistry

Formalin fixed, paraffin sections were cut at 5-6 microns and rehydrated to water. Heat induced epitope retrieval was performed with FLEX TRS High pH Retrieval buffer (pH 9.01; Agilent Technologies, #K8004; Santa Clara, CA) for 20 minutes (Fig. 1A, C and Fig. 3A). After peroxidase blocking, the antibody LGR5 rabbit monoclonal clone STE-1-89-11.5 (Miltenyi Biotec, #130-104-945) was applied at a dilution of 1:50 (Fig. 1A, C; Fig. 7B; Fig. S10) or 1:100 (Fig. 2A) at room temperature for 60 minutes. The FLEX HRP EnVision System (Agilent Technologies) was used for detection with a 10 minute DAB chromagen application. Slides were counterstained with Harris Hematoxylin for 5 seconds and then dehydrated and coverslipped (University of Michigan Comprehensive Cancer Center Tissue Core Research Histology and IHC Laboratory). Note, sections freshly cut were compared to those that were stored at room temperature for 4 weeks, and showed more robust LGR5 staining (data not shown). The colon cancer tissue microarray (TMA) (Fig. 2A; 2 normal, 3 adenoma, 70 adenocarcinoma, 2 cores per specimen) was freshly cut and provided by BioChain Institute, Inc. (Newark, CA; Z7020032, lot B508131). Alternatively, retrieval was performed with R-Buffer B (pH 8.5; Electron Microscopy Sciences; Hatfield, PA) in a pressurized Retriever 2100 (Electron Microscopy Sciences) overnight (Fig. S1A, Fig. 3B). After peroxidase blocking, the antibody was applied at a dilution of 1:1000 at 4°C overnight. ImmPACT DAB kit (Vector Labs, SK-4105; Burlingame, CA) was used, with DAB chromagen applied for 40-45 seconds.

IHC images were taken and scanned on an Aperio AT2 instrument by the University of Michigan Pathology departmental Slide Scanning Service at 40X (Fig. 1A, B top panel, C^1^, C^2^, and D; Fig. 2A; Fig 7B; Fig. S10) and on an IHC Olympus SZX16 at 8x zoom (Fig. 1B bottom panel). 100X insets were captured on a Zeiss Axio M1 (Fig. 1A^2^). Brightfield images were captured on an Olympus IX70, with a DP71 digital camera (Fig. 3A, B).

### LGR5 Immunohistochemistry scoring for staining intensity in the epithelium and in the stroma

The TMA, along with 5 additional FFPE normal colon samples from autopsies, were scored for staining intensity in both the epithelium and then separately in the stroma (Fig. 2). Two FFPE biopsied large (> 10mm) adenomas, used to derive organoids for this study, were also assessed for reactivity (specimen 282 and 590), but these were not simultaneously scored since they were stained under different conditions (Fig. 3C left column). Scoring was conducted by two independent viewers on blinded samples at 8X and 20X magnification. Scoring key: 0 = non-specific or < 1% staining; 1 = 1-10% or only evident at 20X magnification; 2 = 10-50% or light diffuse staining >50%; 5 = >50%. A standard pathology peer review method was used to resolve discrepancies in scoring between two independent scorers, with a third party reviewer intervening when necessary (Morton et al., 2010). The vast majority of samples were in concordance between the two viewers. Stage I (n=24), Stage II (n=24), and Stage III (n=21) tumors were compared to normal colon (n=7) and adenoma (n=3). TMA cancers with grades I and I-II were grouped (termed “Grade I”; n=20) and cancers grade II-III and III were grouped (termed “Grade III”; n=10) for further analyses. LGR5 stromal and epithelial staining for adenomas (n=3), and Grade I, II (n=38), and III cancers were compared to normal colon tissue (n=7). Differences in LGR5 staining in the stroma or epithelium by stage or grade were quantified by linear regression.

### LGR5 Immunofluorescence

Rehydrated sections were retrieved with R-Buffer B (pH 8.5; Electron Microscopy Sciences) in a pressurized Retriever 2100 (Electron Microscopy Sciences) overnight. After 3X PBS washing, slides were blocked for 1 hour with 0.5% Triton X-100 and 5% donkey serum in PBS. Slides were incubated overnight at 4°C with the following antibodies in 0.05% Tween 20 and 5% donkey serum: LGR5 clone 89.11 at 1:500 (Fig. 1C^3^C^4^); anti-OLFM4 antibody at 1:500 (Abcam, #ab85046; Cambridge, MA)(Fig. 8C); anti-defensin5a at 1:500 (Abcam, #ab90802)(Fig. S1A); anti-E-cadherin at 1:500 (BD Biosciences, # 610181, San Jose, CA) (Fig. 8C); and anti-E-cadherin at 1:500 (R&D, #AF748)(Fig. S1A). Slides were washed 3X for 5 minutes each, followed by 1 hour room temperature incubation with the following secondaries (1:1000): LGR5 plus DAPI (Fig. 1C^3^/C^4^; Fig. 2A) and OLFM4 (Fig. 8C) secondaries (biotinylated secondary donkey anti-rabbit; Jackson Immuno Research Labs, #711-065-152; West Grove, PA); defensin5a (Fig. S1A) and E-cadherin secondary (Fig. 8C)(donkey anti-mouse 488; Jackson Immuno Research Labs, #715-545-150); E-cadherin secondary (Fig. S1A) (donkey anti-goat 647; Jackson Immuno Research Labs, #705-605-147). Slides were washed 3X. LGR5 and OLFM4 were amplified with the SK4105 TSA Kit with Alexa 594 tyramide (Invitrogen, #T20935). All slides were mounted with Prolong Gold without DAPI (Molecular Probes, #P36930). Images were taken on a Nikon A1 confocal microscope.

### Single Cell Isolation and by Magnetic-Activated Cell Sorting for LGR5(+) and LGR5(−) cells (Fig. 3)

#### Organoid dissociation into single cells

On the day of sorting, the cultures were treated with 10μM Y27632 for 2.5 hrs prior to harvest. Matrigel was digested for 45 minutes with cold 2mM EDTA in DPBS and organoids were washed 3X with cold DPBS (100xg at 4°C). Structures were dissociated into single cells using the Tumor Dissociation Kit (human) (Miltenyi Biotec, #130-095-929) with the protocol modifications described as follows. The enzymes designated “H” and “R” were prepared in warm HBSS modified to 0.13mM calcium and 0.9mM magnesium [10% HBSS, calcium, magnesium, no phenol red (Gibco, #14025092) with 90% HBSS, no calcium, no magnesium (Gibco, #14170112)] to minimize differentiation of the epithelial cells while supporting enzymatic activity. The organoids were suspended in 20mL of enzyme solution containing 5μM Y27632, then transferred to two BSA-coated C-tubes (Miltenyi Biotec, #130-096-334) and dissociated with a gentleMACS Dissociator (Miltenyi Biotec, #130-093-235) using the program *h_tumor_01*, once at the onset and every 15 minutes for 1 hour, and finishing with two program runs. Between runs the suspension was slowly rotated at 37°C. The cell suspension was then poured over a succession of cold BSA-coated cell strainers into 0.5% BSA-2mM EDTA-PBS: 100μm (Corning, #DL 352360) to 40μm (Corning, #DL 352340) to 20μm (CellTrics of Sysmex Europe GmbH, #04-0042-2315; Norderstedt, Germany). The cells were washed 3X in the 0.5% BSA-2mM EDTA-PBS and centrifuged for 5 minutes at 500xg. Cells were counted under trypan blue exclusion to estimate live cell number using a Countess Automated Cell Counter (Invitrogen, #C10227).

#### Magnetic bead-LGR5 antibody labeling

Single cells were resuspended and processed in cold 0.5% BSA-2mM EDTA-PBS containing 5μM Y27632 in a BSA-coated tube (using BSA-coated tips for mixing) pelleted at 500xg, and otherwise stained according to the product instructions (Miltenyi Biotec: Anti-human LGR5 MicroBeads #130-104-072; clone 22H2.8; APC-Check Reagent #130-098-892).

#### Magnetic separation

Stained cells were resuspended at ≤10^7^ cells/mL in a DNAse Buffer containing HBSS (1:10 Ca^2+^Mg^2+^), 0.5% BSA, 5μM Y27632, 200 Kunitz units/mL DNAse (Sigma-Aldrich, #D5025). Cells were applied over the columns (Miltenyi Biotec, LS Columns, # 130-042-401) through a cold BSA-coated 20μm cell strainer at 1mL per column. Columns were prepped by coating in the DNAse Buffer for 45 minutes at 4°C prior to use. All washes were performed with the DNAse Buffer. After removing the column from the magnet, 2mL DNAse Buffer was applied to the column to flush out the magnetically-bound cells into a 15mL BSA-coated tube. Unlabeled flow-through cells (magnet intact) and labeled magnet-bound cells were pelleted at 500xg for 5 minutes and resuspended in 0.1% BSA-PBS-2mM EDTA with 10μM Y27632.

### Flow Cytometry

Cells were analyzed on a LSRII cytometer (BD Biosciences) and sorted on a MoFlo Astrios (Beckman Coulter; Brea, California) instrument at the University of Michigan Cancer Center Flow Cytometry core facility. Events first passed through a routine light-scatter and doublet discrimination gate, followed by exclusion of dead cells using 4’,6-diamidino-2-phenylindole (1μM DAPI dilactate; Molecular Probes, #D3571). Gating strategy set the APC-positive cell population at 0.05-0.1% of the viable MACS flow-through cells, LGR5(FT+) (Fig. 3C bottom panels; all FACS RNA sequencing-derived data; Fig. 8A, specimen 81, 83). Gating strategy set the APC-positive cell population at 0.05-0.1% of the viable FMO-APC check reagent control for the analysis of samples that were not MACS processed (Fig. 3C top and middle rows; Fig. S2A). For RNA expression analyses, cell fractions were sorted into RLT lysis buffer (Qiagen). Flow cytometric data analysis was performed using Winlist 3D software (Verity) and FlowJo vX.0.7 (Tree Star). For each specimen MACS-processed, three cell fractions were FACS and collected (Fig. 3A, C bottom panel; Fig. 8A; all FACS RNA-sequencing-derived data): MACS magnet-bound positives, LGR5(+); MACS magnet-bound negatives, LGR5(−); and MACS unbound flow-through, LGR5(FT-).

### High Throughput RNA Sequencing (RNA-seq)

RNA was extracted from sorted cells using the RNeasy Micro Kit (Qiagen, #74004) with on-column DNase digestion. RNA concentration and quality were determined using a Nanodrop (Thermo Scientific) and Bioanalyzer (Agilent). Due to the small number of cells following FACS and corresponding low level of input RNA, we depleted ribosomal RNAs with RiboGone (Takara Clontech, #634846; Mountain View, CA) and prepared sequencing libraries utilizing the SMARTer Stranded RNA-Seq kit (Takara Clontech, # 634839) following the manufacturer’s recommended protocol. Libraries were multiplexed over 2 lanes and sequenced using paired end 75 cycle reads (sample 14881) and 125 cycle reads (samples 282, 584, and 590) on a HiSeq sequencer (Illumina) at the University of Michigan DNA Sequencing Core Facility.

### RNA-Seq Data Analysis

#### Alignment and quality control

Computational analyses were performed using the Flux high-performance computer cluster hosted by Advanced Research Computing (ARC) at the University of Michigan. Raw sequencing read quality was assessed utilizing FastQC. The first five nucleotides of the both reads in each read pair were trimmed with SeqTK due to the presence of adapter sequence and high nucleotide redundancy. A splice junction aware build of the human genome (GRCh37) was built using the genomeGenerate function from STAR 2.3.0 (Dobin et al., 2013). Read pairs were aligned to the genome using STAR, using the options “outFilterMultimapNmax 10” and “sjdbScore 2”.

#### Differential expression testing

The aligned reads were assigned to genomic features (GRCh37 genes) using HTSeq-count, with the set mode “union”. We conducted differential expression testing on the assigned read counts per gene utilizing edgeR (Robinson et al., 2010). Analysis were conducted to compare expression of LGR5(+), LGR5(−), and LGR5(FT-) cells, adjusting for study subject as a covariate using glmLRT. To reduce the dispersion of the dataset due to lowly expressed genes, genes with a mean aligned read count less than five across all samples were excluded from analysis. Normalized counts per million were estimated utilizing the “cpm” function in edgeR. Genes were considered differentially expressed between conditions at a false discovery rate adjusted p-value < 0.05 (Fig. 4; Fig. S3; Table S2).(Benjamini and Hochberg, 1995).

**Fig. S3 (related to Fig. 4).**
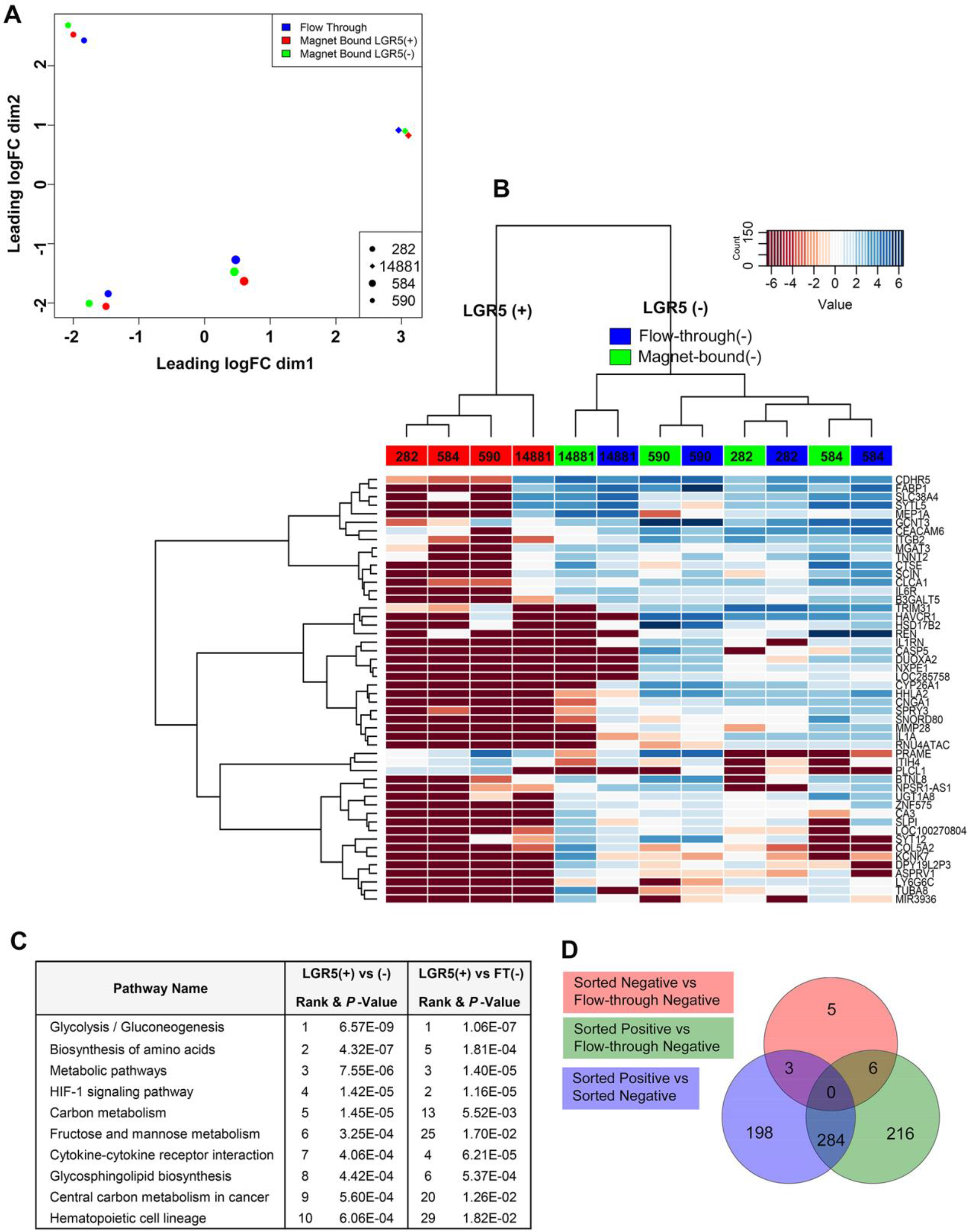
Transcriptomic profiling of human LGR5(+) primary adenoma cells; comparisons between sorted MACS magnet-bound Lgr5(−) and sorted MACS flow-through LGR5(FT-) cells. **(A)** Multidimensional scaling plot of the LGR5(+), sorted MACS magnet-bound Lgr5(−) cells, and sorted MACS flowthrough LGR5(FT-) cells based on the top 500 most variable genes. **(B)** Unsupervised hierarchical clustering heatmap of the 50 most variable genes between the LGR5(+) (red), Lgr5(−) (green), and LGR5(FT-) (blue) populations. **(C)** The top 10 most enriched KEGG pathways for differentially expressed genes between the LGR5(+) and Lgr5(−); and LGR5(+) and LGR5(FT-) cells. **(D)** Gene expression overlap between the [LGR5(+) vs. Lgr5(−)], [LGR5(+) vs. LGR5(FT-)], and [Lgr5(−) vs. LGR5(FT-)].

#### Comparison to previously published datasets

RNA-seq data from normal colon (samples ERR315348, ERR315357, ERR315400, ERR315403, ERR315462, ERR315484 from Tissue Based Map of the Human Proteome Science REF) were processed as described above, and differential gene expression between the LGR5(+) cells and normal colon were calculated as described above (Fig. 6B, C). Overlap between genes overexpressed in LGR5(+) cells with the previously reported *Lgr5*+ ISC gene signature (Munoz et al., 2012) was calculated by identifying the overlap between genes present in both the Muñoz et al ISC gene expression signature and genes identified as upregulated in LGR5(+) adenoma cells (Fig. 6A; Table 2; Table S2). Statistical significance of the overlap between these gene signatures was calculated using the hypergeometric distribution. Genes in the Muñoz et al ISC signature that were not present at detectable levels in our dataset were excluded from this calculation.

#### Pathway analyses

Differentially expressed pathways were identified utilizing iPathways (Advaita; Plymouth, MI). A directional analysis was conducted on all genes by including *P*-value of the differential expression test as a measure of effect size and log2 fold difference in expression as a measure of effect direction. KEGG biological pathways and Gene Ontology biological processes were considered differentially expressed at a *P*-value <0.05 (Fig. S4-9; Table S2).

**Fig. S4 (related to Fig. 4).**
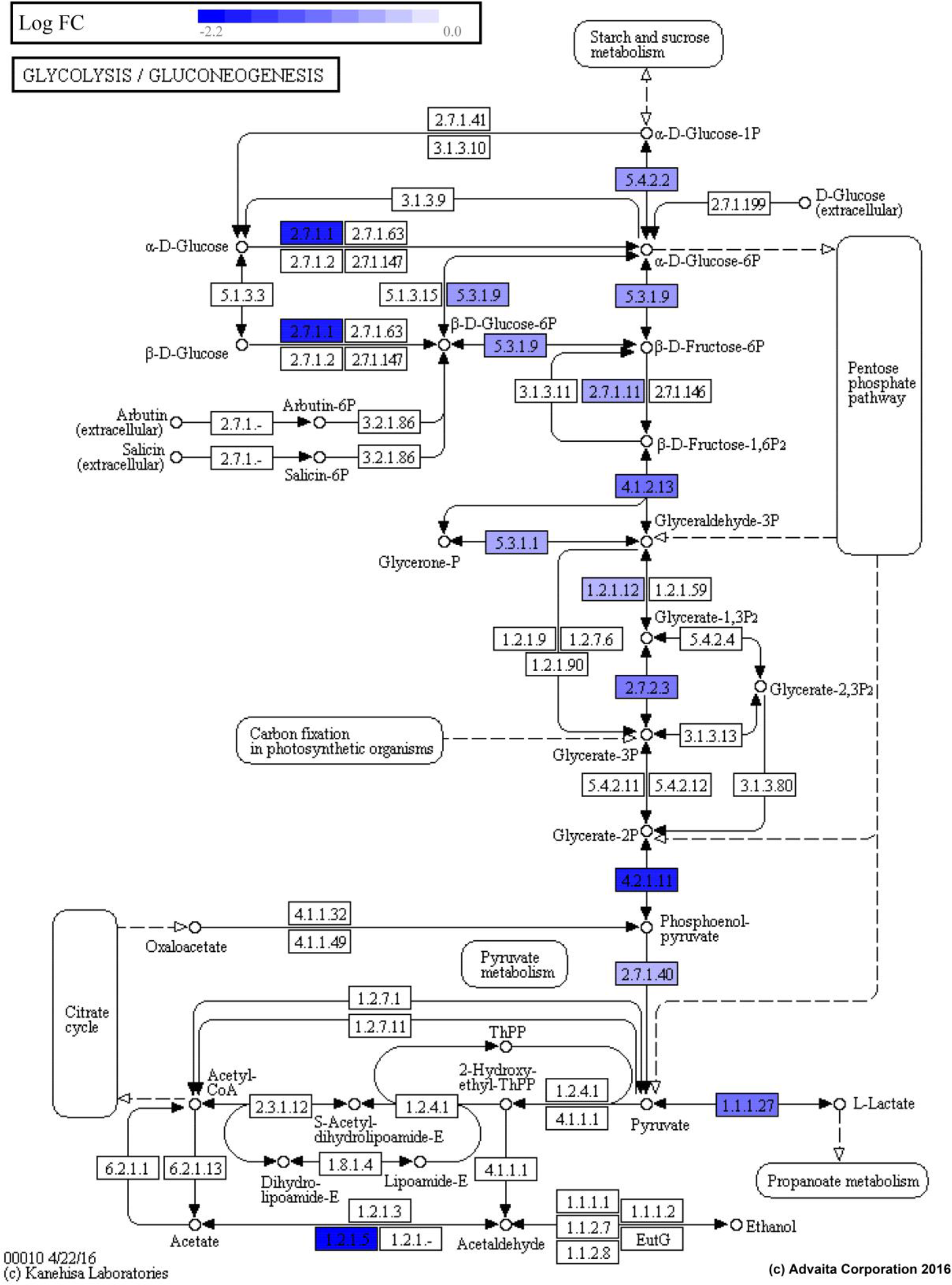
Relative expression of genes involved in the KEGG pathway “Glycolysis and Gluconeogenesis” between LGR5(+) and LGR5(−) human adenoma cells (*P*-value=6.57E-09).

**Fig. S5 (related to Fig. 4).**
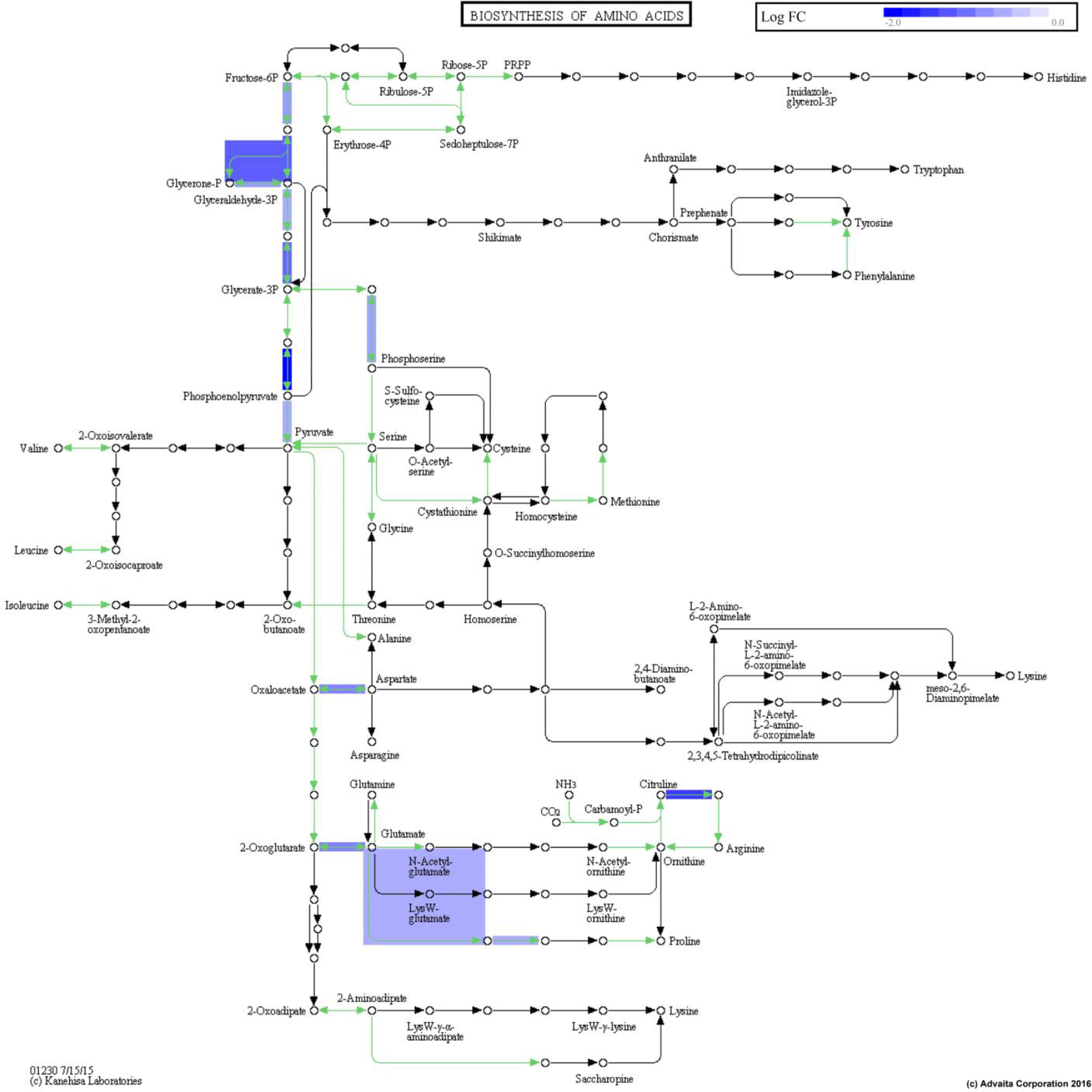
Relative expression of genes involved in the KEGG pathway “Biosynthesis of Amino Acids Pathway” between LGR5(+) and LGR5(−) human adenoma cells (*P*-value=4.32E-07).

**Fig. S6 (related to Fig. 4).**
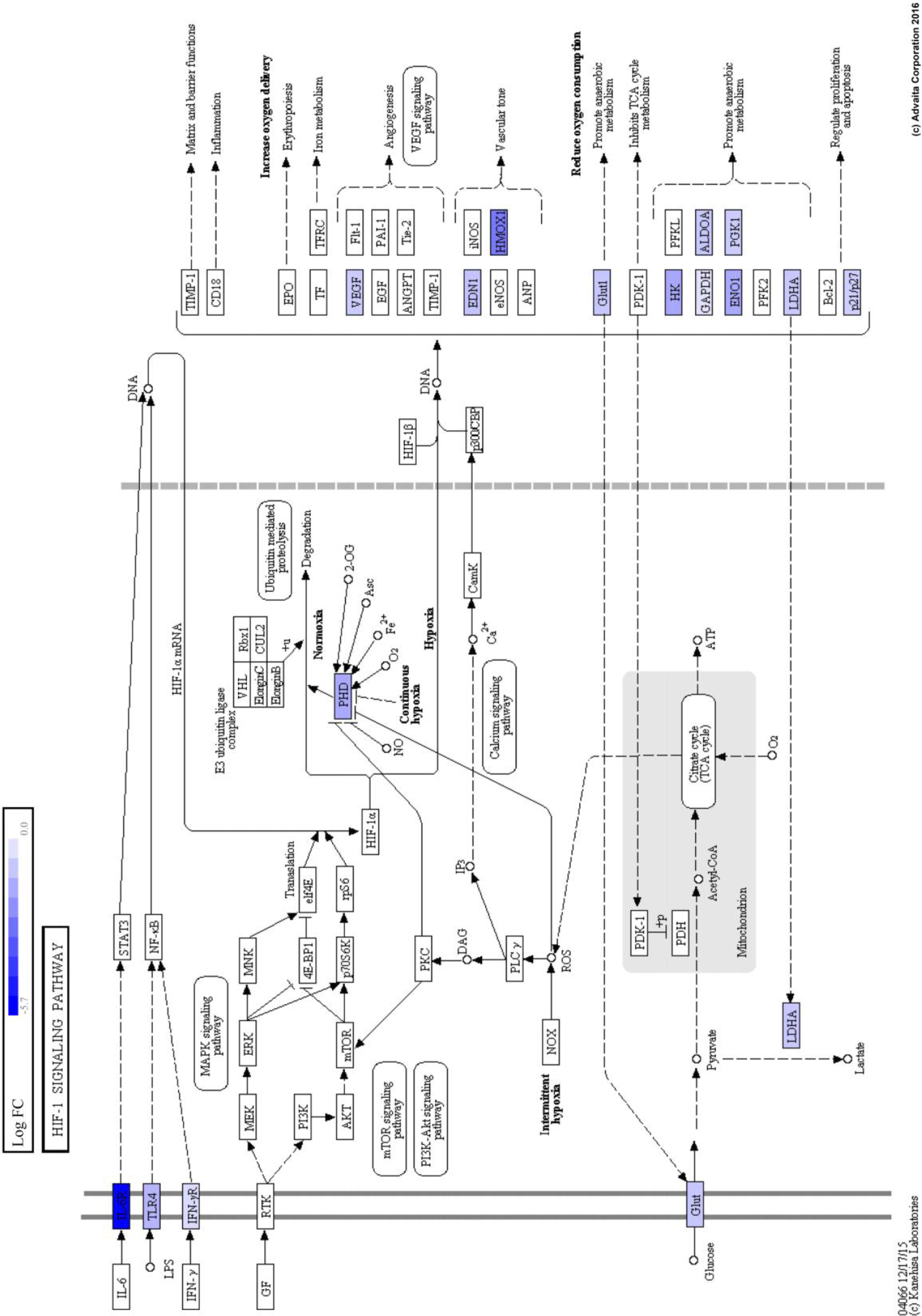
Relative expression of genes involved in the KEGG pathway “HIF-1 Signaling Pathway” between LGR5(+) and LGR5(−) human adenoma cells (*P*-value=1.42E-05).

**Fig. S7 (related to Fig. 4).**
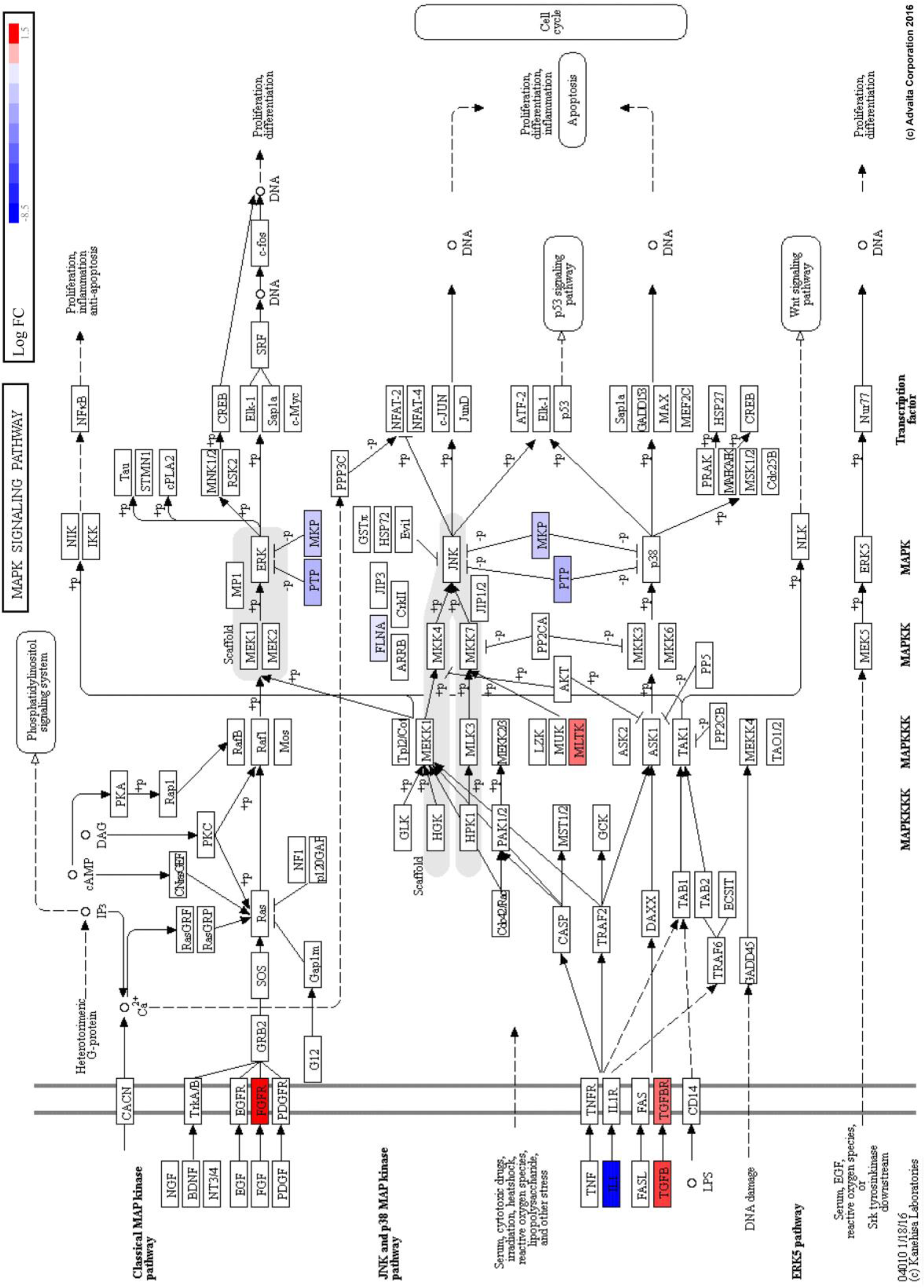
Relative expression of genes involved in the KEGG pathway “MAPK Signaling Pathway” between LGR5(+) and LGR5(−) human adenoma cells (*P*-value=.0105).

**Fig. S8 (related to Fig. 4).**
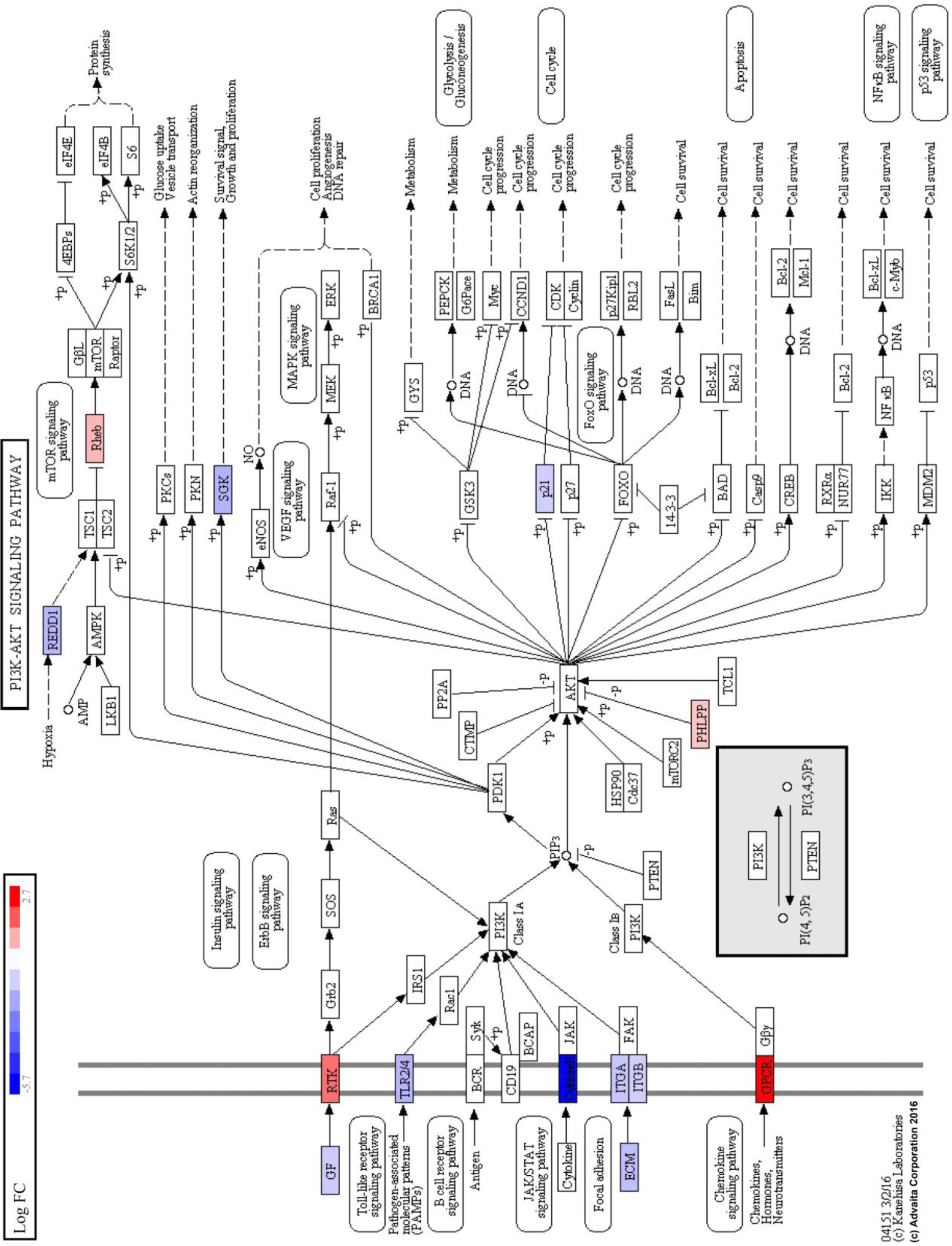
Relative expression of genes involved in the KEGG pathway “PI3K AKT
Signaling Pathway” between LGR5(+) and LGR5(−) human adenoma cells (*P*-value=.0102).

**Fig. S9 (related to Fig. 4).**
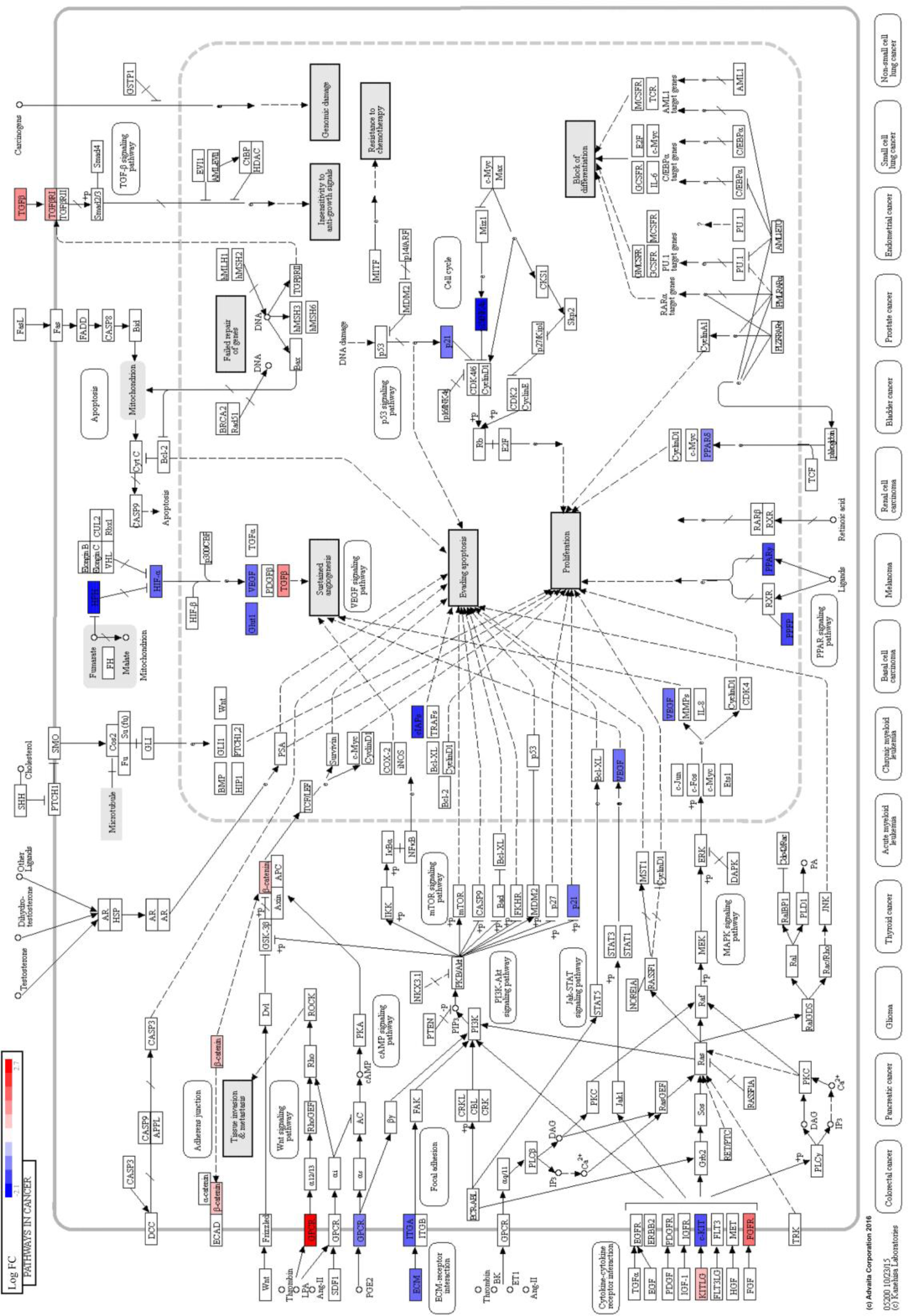
Relative expression of genes involved in the KEGG pathway “Pathways in Cancer” between LGR5(+) and LGR5(−) human adenoma cells (*P*-value=.0133)

**Fig. S10 (related to Fig. 4).**
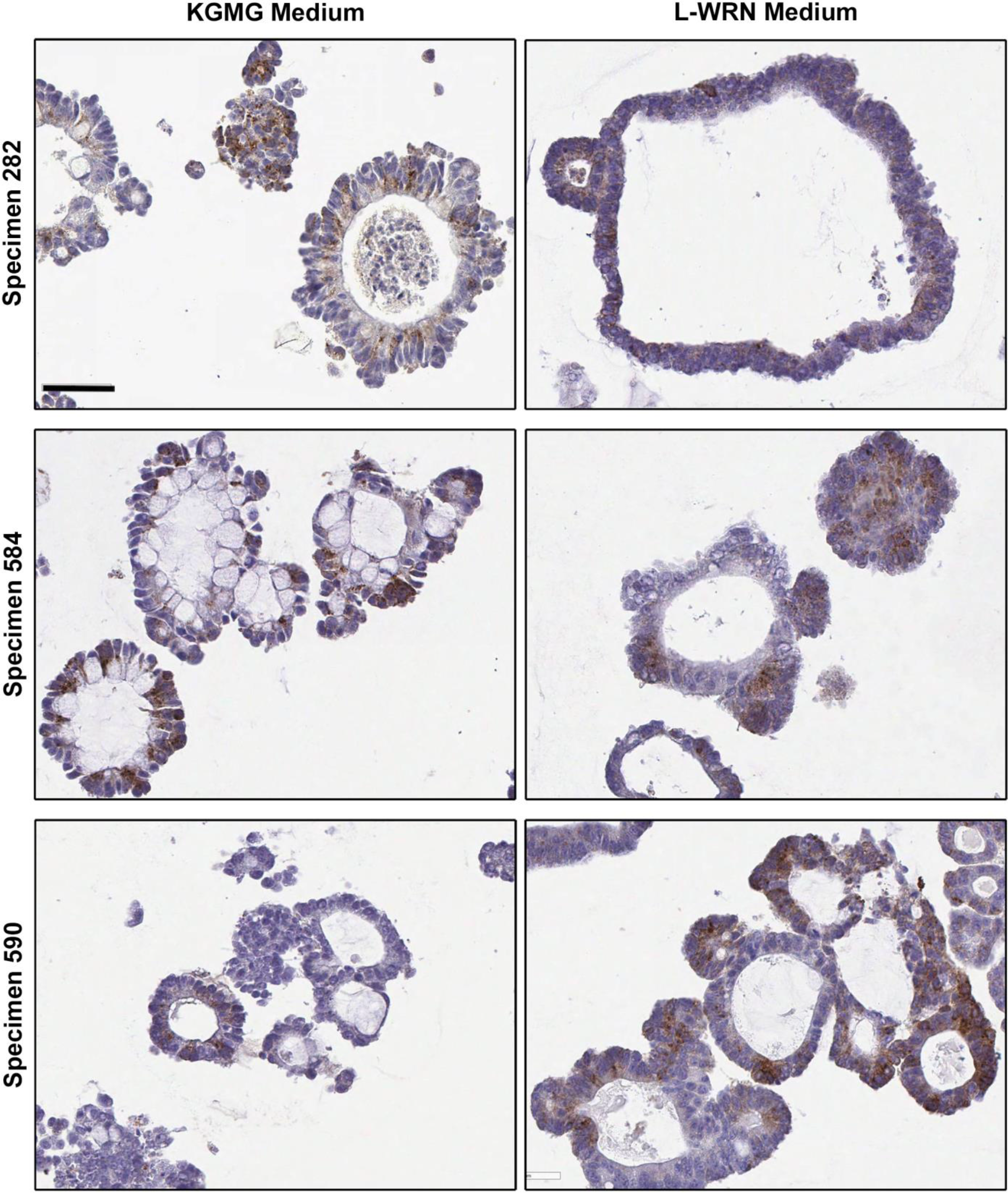
LGR5 expression measured by immunohistochemistry in 3 organoid lines grown in either KGMG or L-WRN media. Scale bar, 50μm.

### Association of SMOC2 and DKK4 expression with LGR5 expression

#### *DKK4* and *LGR5* quantitative real-time PCR of adenoma organoids grown in L-WRN vs. KGMG

Cultured organoids from three adenomas, 282, 584, and 590, were transitioned from KGMG to L-WRN, with the concomitant shift from budding to cystic structures. *LGR5*, *SMOC2*, and *DKK4* mRNA expression were then assessed as previously described above and presented as increased expression in L-WRN relative to KGMG (Fig. 7A). *DKK4* and *SMOC2* primer sequences were as follows: *hDKK4* (Cui et al., 2007), GGAGCTCTGGTCCTGGACTT (forward primer), TCTGGTATTGCAGTCCGTGT (reverse primer); *hSMOC2*, AGAGAGGTTTCTGGGGCG (reverse primer), CTCAGAAGTTCTCGGCGCT (forward primer).

#### mRNA *in situ* hybridization (ISH) for localization of *SMOC2* and *DKK4* relative to LGR5

Formalin fixed, paraffin sections were cut at 5-6 microns and rehydrated to water. *In situ* hybridization was performed using the RNAscope 2.0 HD detection kit [Advanced Cell Diagnostics (ADC), #310035; Newark, CA] according to the standard provided protocol (Fig. 1B, D; Fig 7C, D), and probes were designed by ADC. The human LGR5 probe was designed using the NCBI LGR5 sequence (NCBI Reference Sequence: NM_003667.2) and targets the region between 560-1589 base pairs (HS-LGR5, #311021, lot 15022A). The human SMOC2 probe was designed using the NCBI SMOC2 sequence (NCBI Reference Sequence:NM_1166412.1) and targets the region between 1649-2609 base pairs (HS-SMOC2, #522921, lot 17192B). The human DKK4 probe was designed using the NCBI DKK4 sequence (NCBI Reference Sequence:NM_014420.2) and targets the region between 17-779 base pairs (HS-DKK4, #429371, lot 16244A). All incubations were performed at 40°C in the ADC HybEZ hybridization system oven (#310010). Images were taken and scanned on an Aperio AT2 instrument by the University of Michigan Pathology departmental Slide Scanning Service.

### Transduction of organoids with lentiviral TCF/LEF GFP reporter and ImageStream Analysis

Colon adenoma specimen 282 organoids were transduced with a lentiviral TCF/LEF GFP reporter (Qiagen, CLS-018G) based on Koo et al (Koo et al., 2013) with the major modifications being that 1) organoids were mechanically dissociated in the absence of enzymatic digestions (to avoid single cells), and 2) that incubations were done at 4°C. Organoids were removed from Matrigel and mechanically dissociated into structures between 20-100 μm in diameter with the gentleMACS using programs *h_Tumor_01.01* followed by *m_Lung_01.01*, and then by passing through a BSA-coated 100μm cell strainer. Organoids were transduced with 10TU viral particles/mL in the presence of 8μg/mL polybrene (Sigma-Aldrich), 10μM Y27632, and 2.5μM CHIR99021 (Miltenyi). Organoids were passaged under puromycin selection and transitioned to cystic morphology with L-WRN medium. Cells were processed and stained for LGR5-APC as described above, without MACS enrichment. Single cells were analyzed and imaged with the Amnis ImageStreamX Mark II (EMD Millipore; Kankakee, IL) for brightfield, SSC, and for co-expression of *TCF/LEF*-GFP and LGR5-APC fluorescence (Fig 5B,C). Dead cells were excluded using the Live/Dead Fixable Near-IR Dead Cell Stain Kit (Molecular Probes, #L34975; Eugene, Oregon). Analysis strategy follows: gradient RMS for focus, to intensity of MC 1 (Live-Dead) for viability, to intensity of MC 3 (GFP-FITC or APC for rough positivity, to area channel 2 (fsc) vs. intensity channel 6 (ssc) for sizing to eliminate debris, to finally refined FITC or APC histograms. Samples were analyzed offline in the Amnis IDEAS software platform to generate images at 60x magnification (Fig 5C,D).

### Normal Colinic Crypt LGR5(+) Cell Isolation and Culture

#### Isolation from Tissue, Single Cell Dissociation, Magnetic Separation and Flow Cytometry

Crypts were isolated from three warm autopsy normal colon specimens (6 cm^2^) and processed as previously described (Dame et al., 2014), a modification of Sato et al (Sato et al., 2011a). Crypts were dissociated into single cells, as described above for the adenoma organoids, and a final 1mL of a dense crypt pellet was processed per C-tube. Magnetic separation and flow cytometry (Becton-Dickinson FACSAria III sorter) were as above, with the added gating/analysis for EpCAM expression using a phycoerythrin (PE)- conjugated antibody (2μg /mL buffer/≤ 10^6^ cells; BioLegend, #324205; San Diego, CA) and an EpCAM isotype control (2μg /mL buffer/≤ 10^6^ cells; BioLegend, #400311). EpCAM-PE(+)/DAPI(−) cells were interrogated for LGR5 expression as done above with MAC processed samples (Fig. 8A, specimen 81 and 83) or with rigorous so-called fluorescence-minus-one controls [FMO; DAPI(−), EpCAM(+)] for sorting of pre-magnet cells (Fig. 8A, specimen 84; Fig. 8D, normal organoid-derived single cells).

#### Culture of FACS LGR5(+) and LGR5(−) cells isolated from normal colon organoids

Normal colon organoid cultures were established from warm autopsy resections (Table 1). LGR5(+)/(−) cells were isolated from early passage organoids (P3-P8) and sorted (2100 cells each) into 13mL ice-cold serum-free growth medium further containing 0.25mg/mL Matrigel. The medium composition was roughly based on previous mouse small intestine/colon single cell organoid-forming efficiency studies(Yin et al., 2014, von Furstenberg et al., 2014, Wang et al., 2013, Shimokawa et al., 2017, Jung et al., 2015). The medium is serum-free, with Wnt3a, R-spondin-3 and Noggin provided by 50% L-WRN cell 24hr-conditioned media [conditioned media: AdvDMEM, 2mM GlutaMax, 10mM HEPES, N2, B27, 1mM N-Acetyl-L-cysteine, 13 μg/mL gentamicin, 25μg/mL Primocin, 5ng/mL EGF, 5ng/mL FGF-2 (Miltenyi Biotec; 130-093-837), and 300ng/mL human Afamin (Mihara et al., 2016) (Sino Biological Inc.; 13231-H08H) to stabilize Wnt3a in the absence of serum]. This was made to 100% complete growth medium with final inclusion of: AdvDMEM, 2mM GlutaMax, 10mM HEPES, N2, B27, 1mM N-Acetyl-L-cysteine, 25μg/mL gentamicin, 50μg/mL Primocin, 100ng/mL EGF, 10μM Y27632, 10mM Nicotinamide, 3μM SB202190, 500nM A83-01, 10nM PGE2 (Sigma-Aldrich; P6532), and 10nM Gastrin (Sigma-Aldrich, G9020). For the first 48hrs, 5μM CHIR99021 and 2μM Jagged-1 (AnaSpec; AS-61298) were also added. Sorted cells were centrifuged at 4°C for 10 minutes at 500xg, and resuspended in the residual media (25uL) and Matrigel (112μL) for 4 wells at 500 cells/28μL Matrigel/well of a pre-warmed 24-well plate. After 10 minutes, 400μL of single cell growth medium was added. On day 8, the medium was changed to L-WRN medium (10% FBS from L-cell conditioned medium), with the added growth factors listed here. Preliminary studies indicated that early inclusion of serum was detrimental to the initiation of growth (data not shown). At day14-17, images were taken and organoids over 100μm were counted to access organoid-forming efficiency (Fig. 8D). Cultures were passaged for approximately 2 months with cryopreservation before ending.

**Table S1 (Related to Results):** Enrichment of LGR5(+) cells by magnetically activated cell sorting as assessed by FACS. Cells were derived from adenoma organoids cultured in L-WRN medium. Each analysis was from a separate experiment. Percentages displayed represent mean values averaged across experiments.

| Specimen | No magnet<br>(LGR5+% of all cells) |  | Magnetically Activated Cell Sorting (MACS)<br>(LGR5+% of Magnet-bound Cells) |  | MACS<br>Fold<br>Enrichment |
| --- | --- | --- | --- | --- | --- |
| 590 | 12.90% | n=1 | 5.52% | n=3 | 0.43 |
| 282 | 0.24% | n=1 | 7.04% | n=2 | 29.33 |
| 584 | 1.51% | n=1 | 6.43% | n=2 | 4.25 |
| Average |  |  |  |  | 11.34 |
| Standard Error |  |  |  |  | 6.55 |

**Table S2 (Related to Fig. 4-6 and Fig. S3, 4-9):**
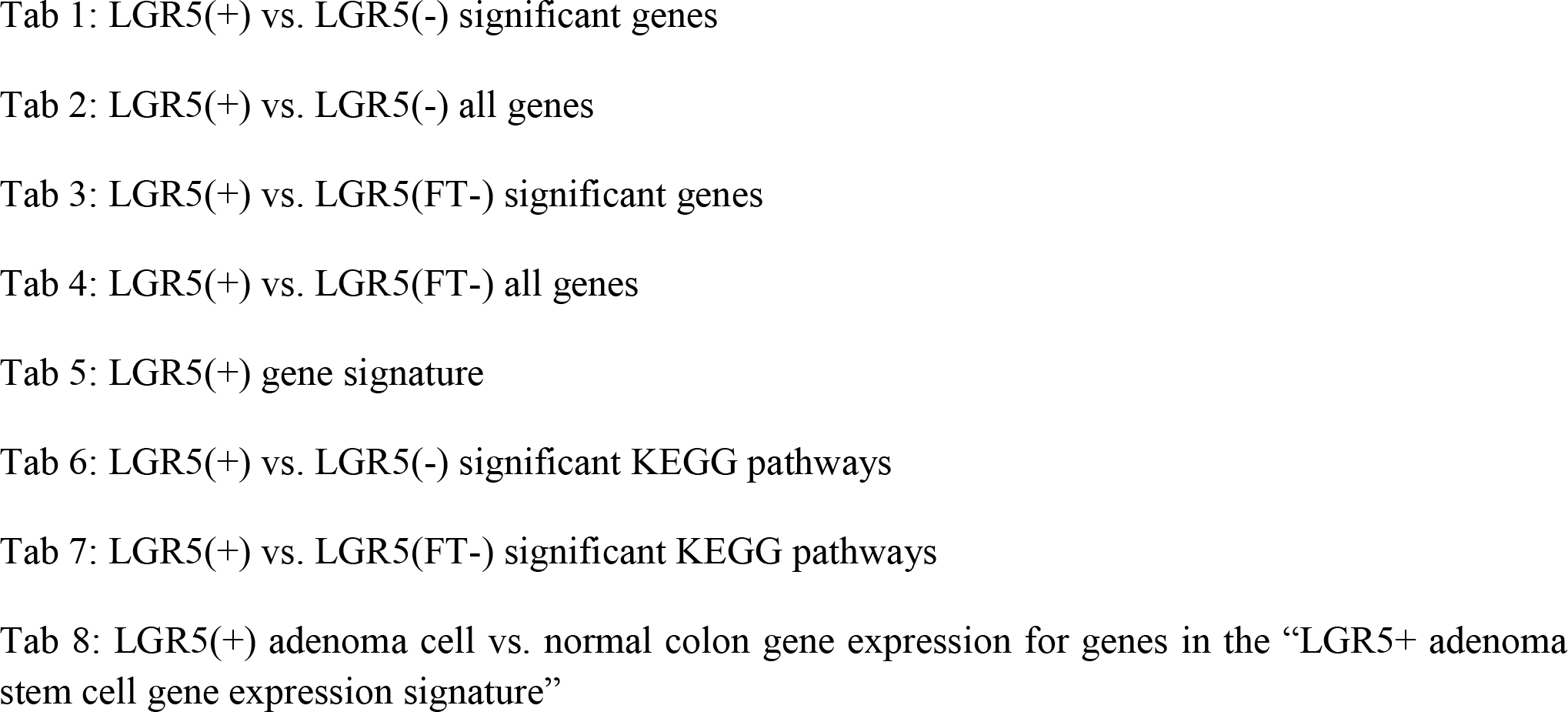
LGR5(+) Adenoma RNA-seq analyses

